# Identification of Low-Complexity Domains by Compositional Signatures Reveals Class-Specific Frequencies and Functions Across Reference Proteomes

**DOI:** 10.1101/2023.07.24.550260

**Authors:** Sean M. Cascarina, Eric D. Ross

## Abstract

Low-complexity domains (LCDs) in proteins are typically enriched in one or two predominant amino acids. As a result, LCDs often exhibit unusual structural/biophysical tendencies and can occupy functional niches. However, for each organism, protein sequences must be compatible with intracellular biomolecules and physicochemical environment, both of which vary from organism to organism. This raises the possibility that LCDs may occupy sequence spaces in select organisms that are otherwise prohibited in most organisms. Here, we report a comprehensive survey of LCDs in all known reference proteomes (>21k organisms), with added focus on rare and unusual types of LCDs. LCDs were sorted into a rich hierarchical database according to both the primary amino acid and secondary amino acid in each LCD sequence, facilitating detailed comparisons of LCD class frequencies across organisms. Examination of LCD classes at different depths (i.e., domain of life, organism, protein, and per-residue levels) reveals unique facets of LCD frequencies and functions. To our surprise, all 400 LCD classes occur in nature, although some are exceptionally rare. A number of rare classes can be defined for each domain of life, with many LCD classes appearing to be eukaryote-specific. Multiple eukaryote-specific LCD classes could be linked to consistent sets of functions across organisms. Our analysis methods enable simultaneous, direct comparison of all LCD classes between individual organisms, resulting in a proteome-scale view of differences in LCD frequencies and functions. Together, these results highlight the remarkable diversity and functional specificity of LCDs across all known life forms.

## Introduction

In known organisms, the majority of proteome space is devoted to “high-complexity” regions comprised of a diverse mixture of amino acids. Nearly all organisms also contain “low-complexity” domains (LCDs), which have highly skewed amino acid compositions often favoring one or two amino acids [1]. As such, LCDs are a heterogeneous collection of domains with highly divergent biochemical and biophysical properties that depend upon which specific amino acid(s) are enriched in each domain. For example, LCDs enriched in polar and charged amino acids are often intrinsically disordered, whereas LCDs enriched in other amino acids exhibit a greater tendency to adopt ordered conformations [2,3]. Classification of LCDs based on their primary and secondary amino acids (i.e. the most common and second-most common amino acids, respectively, within the LCD) results in a hierarchy of LCD categories that correlate with specific molecular functions [4]. Consequently, such classification schemes are vital for properly interpreting statistical associations between LCDs and molecular functions. Statistics performed on LCDs as a single category will be heavily skewed by the predominant LCD classes in a given dataset, so conflation of distinct LCD categories can lead to overgeneralized and imprecise conclusions.

LCDs from each class differ in their abundance within and across organisms [4,5]. One plausible explanation for these differences is that particular classes of LCDs are tolerated or beneficial to varying degrees in different organisms. A variety of LCDs from multiple LCD classes undergo liquid-liquid phase separation *in vitro* and enable recruitment to biomolecular condensates *in vivo* in response to changes in their surrounding environment (for review, see [6–11]). These changes include shifts in simple physicochemical properties such as temperature, pH, salt concentration, osmolarity, and pressure, as well as large-scale alterations in the abundance of free biomolecules such as RNA, ATP, polyphosphate, polyamines, and polyADP-ribose. In some cases, LCDs act as direct sensors of physicochemical environment in various contexts and in a manner specific to both the LCD type and the change in environment. For example, S/T-rich LCDs were recently linked to CO_2_ sensing in *Candida albicans* [12], M-rich LCDs have been associated with redox sensing in *Saccharomyces cerevisiae* [13,14], Q/H-rich LCDs have been associated with pH sensing in *Drosophila melanogaster* [15,16], and Q-rich LCDs have been linked to temperature sensing in *Arabidopsis thaliana* [17].

While some environmental conditions are buffered by non-equilibrium, physiological regulatory systems, the full complement of known species spans a broad range of ecological niches. Such a diverse collection of niches likely necessitates proteome-scale adaptations in the organisms that occupy them. Given the remarkable responsiveness and environmental sensitivity of LCDs, we reasoned that at least one of the proteome-scale adaptations could be the overall LCD content profile for each organism, including both the abundance of LCDs from each LCD class and the features of those LCDs. Certain LCDs that might be maladaptive to a particular organism might be benign or even functional to a different organism in a different context. Additionally, specific organisms may develop concomitant proteome adaptations that influence LCD tolerability. For instance, specific adaptations in the proteostasis machinery of the model slime mold, *Dictyostelium discoideum*, and of the malarial parasite, *Plasmodium falciparum*, suppress the *in vivo* aggregation of N-rich LCDs [18–21], which are notoriously aggregation-prone and are remarkably abundant in their proteomes. Similarly, K/Y-rich and Y/W/G-rich LCDs in secreted mussel adhesion proteins depend upon post-translational modification of Y residues for their adhesive functions [22,23]. This process is sensitive to disruption by oxidative damage, but this damage is buffered by co-secreted proteins [24].

Broadly, these examples highlight: 1) LCDs that would likely be deleterious in many organisms can have beneficial functions in specific organisms and contexts, and 2) co-adaptations in the proteomes of these organisms enable the functions of these LCDs and/or mitigate their potential negative consequences.

There are many outstanding questions that arise from this line of reasoning. Are all types of LCDs observed in nature? What are the limits of LCD content that are compatible with known life forms? Are rare LCDs endowed with specialized functions in particular sets of organisms?

To begin to address these questions, we performed a comprehensive survey of LCDs in a comprehensive set of reference proteomes. Our method of categorizing LCDs enables detailed quantification of LCD abundance for each LCD class and direct comparisons of whole-proteome LCD content across organisms for each LCD class. We explore rare and unusual LCDs at multiple levels (i.e., per-organism, per-protein, and per-residue levels) and discuss the unique insights offered by each layer, including functions associated with specific classes of LCDs.

## Results

### Measures of LCD Categorization and LCD Rarity

LCDs can be categorized based on the most-enriched amino acid in each LCD sequence, resulting in 20 “primary” LCD classes (one for each of the 20 canonical amino acids). Primary LCD classes differ in their frequencies and abundances across organisms [4], which is likely due to factors including (but not limited to) whole-proteome amino acid frequencies and/or specialized LCD functions. Additionally, primary LCD classes can be further decomposed into “secondary” LCD classes defined by a second amino acid that is co-enriched in the LCD sequence, resulting in an LCD classification hierarchy that correlates with functional specificity [4]. In order to identify instances of primary and secondary LCD classes, an updated set of UniProt reference proteomes was evaluated using the *LCD Compo*sition *S*cann*er* (LCD- Composer [4,25]; see Methods). By convention, primary LCD classes are here defined as regions ≥20 amino acids in length with ≥40% of the composition corresponding to a single type of amino acid. LCDs can be further classified into “secondary” LCD classes, here defined as regions with ≥40% of the composition corresponding to a single “primary” amino acid and ≥20% of the composition corresponding to a “secondary” amino acid (e.g. S-rich LCDs with a secondary bias for R would be instances of the “SR” secondary LCD class). Using this classification scheme, LCD “rarity” was then examined on a variety of related levels (discussed in separate sections below), each with unique implications and insights. Complete datasets of primary and secondary LCDs for all UniProt reference proteomes are available in a publicly accessible repository [26].

### Organism-level LCD Class Frequencies

We first examined what percentage of organisms contain at least one instance of each class of LCDs, which we term “organism-level LCD frequency”. For most primary LCD classes, a moderate to high percentage of organisms contain at least one protein with an LCD of that class (Fig 1, Fig S1, and Tables S1-S5). Consistent with our previous observations [4], organism-level LCD frequency tended to increase in the order of Viruses◊Archaea◊Bacteria◊Eukaryota, with many viruses often lacking an LCD, most likely due to small proteomes. Interestingly, ≥90% of all eukaryotes contain at least one instance for each primary LCD class except W-rich LCDs, which were rare across all 4 domains of life. W is typically rare in proteomes, which likely contributes to the observed rarity: however, other LCD classes represented by rare amino acids (e.g. C, M, and H) are much more common among eukaryotes and occur in other domains of life, potentially suggesting that W-rich LCDs are uniquely limited by selection or lack beneficial activities. Remarkably, none of the 380 secondary LCD classes are completely absent from nature (Fig S1), though some are exceedingly rare (e.g., only 6 total WM LCDs were identified) and likely require experimental validation.

**Fig 1.**
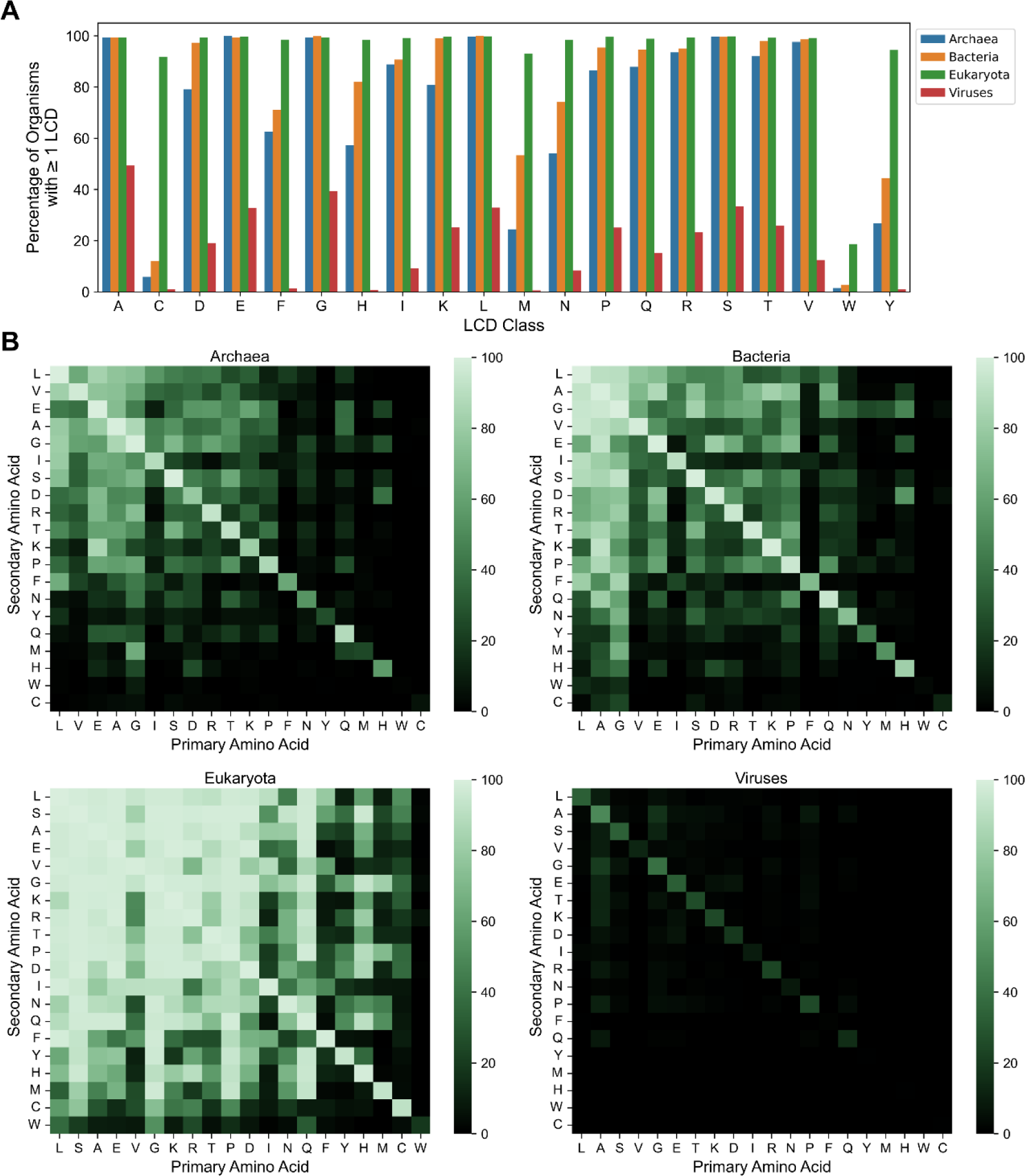
Organism-level LCD frequencies for primary and secondary LCD classes among each domain of life. (A) Percentage of organisms with ≥1 LCD instance for each of the 20 classes of primary LCDs across the four major domains of life. (B) Heatmaps indicating the percentage of organisms from each domain of life with ≥1 instance of each secondary LCD class. In each plot, the x-axis represents the primary amino acid enriched in the LCD (≥40% composition) and the y-axis represents the secondary amino acid enriched in the LCD (≥20% composition). Values for primary LCD classes are indicated in the diagonals to facilitate relative comparisons. Amino acids in panel B were sorted by average whole-proteome frequency rank.

Each of the major domains of life exhibits similar trends with respect to the percentage of organisms containing at least one instance of each two-residue LCD class, a measure that we will refer to as LCD class representation (Fig 1B and Tables S2-S5). Viruses exhibit universally low LCD class representation. For the remaining domains of life, there are clear islands of well- represented LCD classes separated by channels of poorly represented LCD classes. These rare LCD classes tend to occur for amino acids that have low whole-proteome frequencies, such as the aromatic amino acids, C, and M. However, there are noteworthy exceptions. First, some amino acids that do not have low whole-proteome frequencies (e.g., V or I) exhibit relatively low LCD class representation. Second, some amino acids that have low-whole-proteome frequencies (e.g., Q and H in eukaryotes) exhibit relatively high LCD class representation. Third, some rare primary LCD classes exhibit specifically higher representation among subclasses of secondary amino acids. For both Y-rich and M-rich primary LCD classes, G is the most common secondary residue in eukaryotes; these YG and MG classes are also detected to a lesser extent in archaea and bacteria. It is worth noting that our method intentionally does not impose an upper-bound restriction on the degree of enrichment of the secondary amino acid, only minimum requirements for both the primary and secondary amino acids. This may result in some degree of overlap in reciprocal secondary LCD classes (e.g., GM and MG classes).

Therefore, while these LCD classes may include LCDs with ≥40% Y or M and still higher G content, these rare amino acids are nevertheless enriched to an unusual degree, and this is very specific to a particular subcategory of LCDs.

### Rare LCDs Have Specific Secondary-Class Preferences and Can Be Linked to Known and New Functional Classes in Well-Studied Organisms

The primary class of C-rich LCDs was relatively rare among archaea, bacteria, and viruses but was quite common among eukaryotes (Fig 1A). Further examination of secondary LCD classes among the C-rich LCDs indicates disproportionate sorting of C-rich primary LCDs into specific secondary LCD classes (Fig 1B), potentially reflecting functional specialization among certain C-rich secondary LCD classes. Therefore, we performed separate GO term analyses in a set of 13 model eukaryotic organisms for all classes with C as the primary or secondary amino acid (i.e., CX or XC classes), with a threshold of ≥5 LCD-containing proteins as the minimum sample size to perform the analysis. Keratin and keratin-related functions were significantly enriched among multiple C-rich LCD classes (predominantly the CS/SC, CT/TC, CG/GC, CP/PC, and CV classes) across many of the model eukaryotic organisms (supplementary data available at [26]). This is both consistent with the known compositional bias of keratin proteins and partially explains why C-rich LCDs are more frequently observed in eukaryotes compared to other domains of life.

Coarse-grained analysis of whole-protein amino acid composition suggested that keratin proteins with distinct biological purposes and keratin proteins from different organisms exhibit unique compositional profiles [27]. The hierarchical classification of LCDs afforded by LCD- Composer enables finer resolution of the LCD regions predominantly responsible for the C bias and enables spatial resolution of distinct C-rich secondary LCD classes within single proteins.

To determine the degree to which the C-rich keratin-related proteins overlapped across LCD classes within each organism, proteins that were annotated with the terms “keratin filament” or “keratinization” were collected for all of the C-rich LCD classes examined by GO-term analysis (Table S6). LCD types and frequencies among keratin-associated proteins differed substantially across organisms (Fig 2). For example, human keratin-associated proteins contained the most diversified set of LCDs, while mouse and rat keratin-associated proteins were more strongly skewed toward CS/SC or CP/PC LCDs, and cow keratin-associated proteins tended to contain a mixture of CS/SC, CP/PC, and CT LCDs. Additionally, the types of LCDs occurring in the same proteins (“co-occurring” LCDs) varied across organisms. For example, the human CG/GC and CV classes exclusively co-occurred with the CS/SC and CP/PC classes, whereas the human CT/TC class co-occurred with the CQ and CR classes in addition to the CS/SC and CP/PC classes.

**Fig 2.**
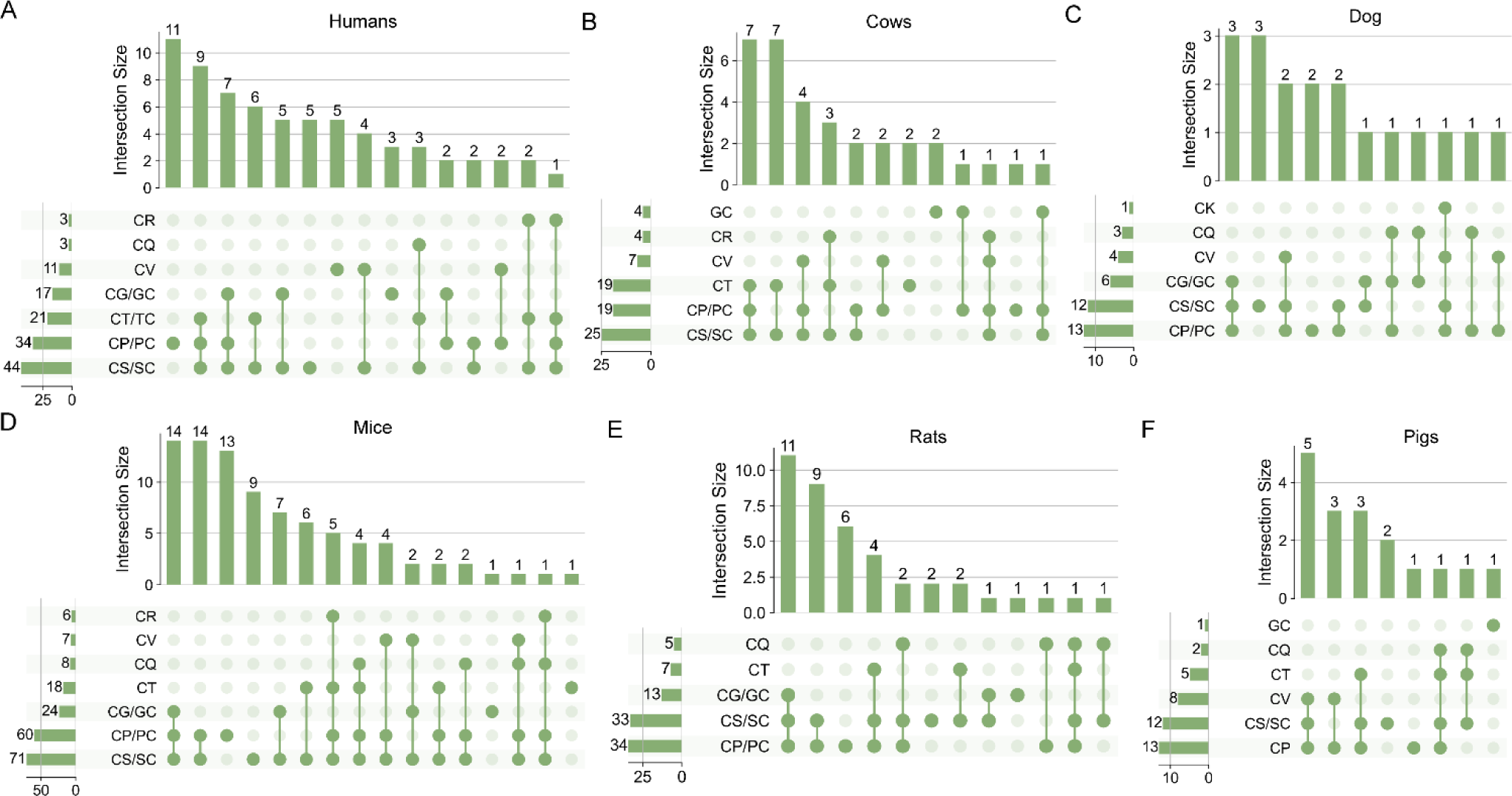
Protein set overlap among keratin-associated proteins identified across distinct LCD classes in model eukaryotes. Keratin-associated proteins with C-rich LCDs were parsed into secondary LCD classes and evaluated for co-occurrence (i.e., two LCD types appearing in the same protein) across LCD classes. Each pair of reciprocal LCD classes (e.g., CS and SC) was grouped into a single representative category. The co-occurrence of LCDs for each organism are depicted in UpSet plots (analogous to a Venn diagram). The graphs on the left of each panel indicate the number of keratin proteins containing each secondary class of C-rich LCDs. The bar graph at the top of each panel indicates the number of proteins with each combination of C-rich secondary LCD classes (which are indicated by green dots and connecting lines below the bar graph). For example, in humans, three keratin proteins contain CR LCDs: two of these proteins also contain CG/GC and CS/SC LCDs, while one contains CG/GC, CP/PC, and CS/SC LCDs.

Given the constraints used to define secondary LCD classes (≥40% of a primary AA and ≥20% of a secondary AA), it is possible that the same LCD sequence could simply be assigned to multiple LCD classes. However, barring a few exceptions, the LCDs of different classes occupied spatially distinct regions in each keratin-associated protein for all organisms (Figs S2- S4). Therefore, not only are C-rich LCDs a largely eukaryote-specific class with corresponding eukaryotic functions, but these C-rich LCDs can be spatially resolved within individual proteins by defining compositional signatures.

While the keratin proteins serve as a robust, prototypical model of C-rich LCDs, a variety of non-keratin functions are also detected for specific secondary LCD-classes in particular organisms. For example, CK LCDs preferentially occur among metallothionein proteins and are significantly associated with functions related to metal ion homeostasis in humans, mice, and dogs. These metallothioneins predominantly bind divalent metals such as zinc, copper, and cadmium via their C residues [28], though their K residues may also play an auxiliary role in metal binding [29,30], structural stability [31], subcellular localization [32], and regulation of steady state metallothionein levels [33]. Although the CG class was associated with keratins in mammals, this class was instead strongly associated with spermatogenesis related functions in fruit flies. The QC LCD class in roundworms (*C. elegans*) was associated with functions related to the unfolded protein response and general stress responses in the ER, as well as pharynx development. Indeed, all of these proteins are members of the *abu* family of proteins, which were identified as ER-localized transmembrane proteins that are upregulated in response to ER stress and improved viability during stress [34]. Based on their homology with the CED-1 mammalian ER receptor protein, it was proposed that the *abu* proteins likely act as receptors for damaged macromolecules, including oxidized lipoproteins: it is tempting to speculate that these domains could play a role in sensing or responding to reactive oxygen species – which are generated during protein folding in the ER – or oxidized proteins. Additionally, these *abu* proteins are upregulated during pharyngeal cuticle development and are secreted into the pharyngeal cuticle, where they have been proposed to play a role in cuticle formation and maintenance via phase separation mediated, at least in-part, by C residues in the QC domains (along with other C-rich cuticle proteins) [35].

Gene duplication has been proposed to play a major role in some protein families, including keratin proteins [36], which might contribute both to the prevalence of certain types of LCDs as well as the functions associated with the LCD-containing proteins. Indeed, when Pfam clan annotations are mapped to LCD-containing proteins from each class, a high percentage of proteins from C-rich LCD classes are associated with a single Pfam clan specifically in eukaryotes (Fig S5). However, low percentages of proteins from nearly all other LCD classes are observed across organisms, suggesting that, in general, the prevalence of most types of LCDs is not due predominantly to gene duplication among specific protein families.

In summary, examination of organism-level LCD class frequencies can be used to identify rare LCD classes, reveal specific preferences for secondary amino acids among rare LCD classes, define eukaryote-specific LCD classes (and potentially LCD classes specific to other domains of life), infer functions of multiple rare LCD classes, and spatially resolve similar types of secondary LCDs within families of related proteins.

### Organisms Containing a Large Number of Proteins with Rare and Eukaryote-Specific LCDs Highlight Widespread LCD-Associated Functions

In light of the functional specificity observed among C-rich LCDs in eukaryotes, we next explored whether other eukaryote-specific LCD classes were represented in high numbers across a large number of organisms. Eukaryote-specific LCD classes were arbitrarily defined as those found in >15% of eukaryotic proteomes but <2% of the proteomes from archaea, bacteria, and viruses (each evaluated independently). Among these classes, HQ LCDs were the most common in eukaryotes, with >800 organisms containing ≥10 proteins with HQ LCDs (Fig 3A and Table S1). Importantly, HQ LCDs also occupy a larger percentage of the proteome, on average, in eukaryotes (0.0048%) compared to archaea, bacteria, and viruses (0.000034%, 0.000036%, and 0.00040%, respectively), indicating that HQ LCDs are not more common in eukaryotes due simply to their larger proteomes. NH LCDs is the next most common with ∼150 organisms having 10 or more LCD-containing proteins, followed closely by the previously examined CV, CS, VC, and CP LCDs. Finally, >50 eukaryotic organisms also had 10 or more proteins from the HL and NM LCD classes.

**Fig 3.**
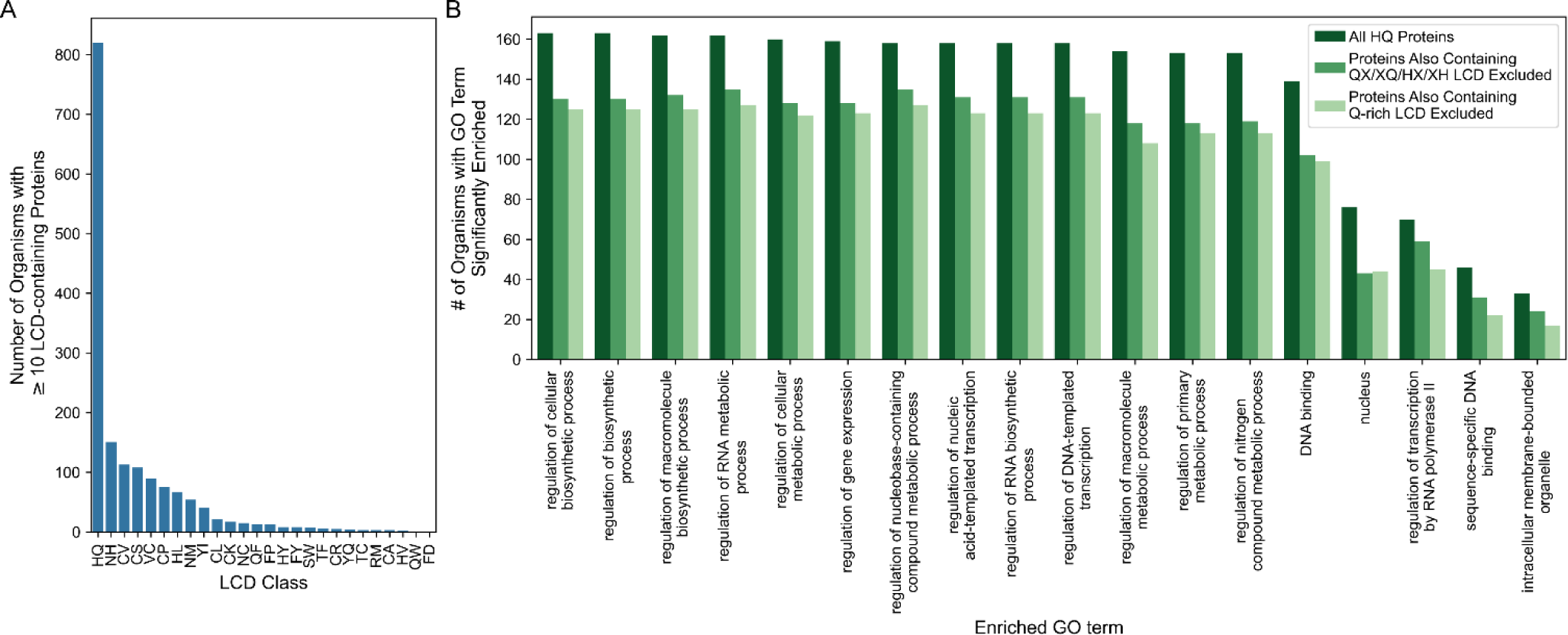
Eukaryote-specific LCD classes can be found in many organisms and exhibit consistent functional associations. (A) The number of organisms with ≥10 LCD-containing proteins for each eukaryote-specific LCD class. (B) Frequency of significant enrichment across organisms for each GO term associated with proteins containing HQ LCDs. GO-term analyses were also performed on the same set of proteins but with those that also contained a spatially distinct QX/XQ LCD (where X is any residue other than H) or a Q-rich LCD (primary class) removed prior to analysis. None of the HQ-containing proteins also had a spatially distinct H-rich LCD (primary class). For simplicity, only GO terms that were significantly enriched for ≥15 organisms and had a minimum depth of 4 in the gene ontology are shown.

In a previous study of 13 metazoan proteomes, Q/H-rich LCDs were found to be significantly associated with transcription-related functions in a small subset of representative organisms [5]. However, the prevalence of HQ LCDs across eukaryotic proteomes potentially indicates a common function linked to HQ LCDs that is widely exploited by many eukaryotes. Additionally, although QH and HQ LCD classes are similar, the compositional constraints imposed during the LCD search can lead to substantially different LCD regions and LCD-containing proteins identified, which in turn could affect downstream functional analyses. We performed GO term analyses for 543 organisms that contained ≥10 HQ proteins and had available gene ontologies. A variety of identical transcription-related terms were significantly enriched across a large fraction of the organisms evaluated (Fig 3B and Table S7), suggesting that the association between HQ LCDs and transcription-related functions is widespread among eukaryotes. Most functions were retained when spatially distinct Q-rich primary LCDs or QX/XQ secondary LCDs (where X represents residues other than H) were removed. None of the proteins contained H-rich primary LCDs that were spatially distinct from the HQ domains.

We next explored whether other LCD classes relatively common among eukaryotes (>50 organisms with 10 or more LCD-containing proteins) could also be linked to specific functions.

The cysteine-rich LCD classes were already explored in the previous section, leaving the NH, HL, and NM LCD classes. GO-term analyses were performed for organisms with available gene ontologies and a set of ≥10 LCD-containing proteins associated with one of these LCD classes (Table S8). All three LCD classes were significantly associated with transcription-related functions across multiple organisms. However, additional functional associations were observed in particular organisms. For example, proteins with NM LCDs were significantly associated with clathrin-related terms specifically in *D. discoideum*. A broader diversity of functions for proteins with NH LCDs were observed and appeared to be organism-specific. For example, proteins with NH LCDs were significantly associated with ubiquitin-protein transferase activity in *Tetranychus urticae*, spermine/spermidine biosynthesis in *Aedes albopictus*, cell morphogenesis in *Arachis hypogaea*, and histone H3 methyltransferase activity in *Meloidogyne enterolobii*.

Together, these examples highlight both functional overlap and functional specificity that can be observed across secondary LCD classes, as well as identify eukaryote-specific LCD classes.

### Protein-level LCD Class Frequencies: LCDs Are More Common than Expected from Whole-Proteome Amino Acid Frequencies and Exhibit Domain-Specific Enrichment of LCD Classes

Organism-level LCD frequencies indicate which organisms contain at least one instance of an LCD but do not indicate how prevalent the LCD is within each proteome. At the protein level, LCD frequency can be defined as the number or percentage of proteins containing an LCD within a single organism. However, key factors influencing protein-level LCD frequencies are whole-proteome amino acid frequencies and proteome size, both of which can vary substantially between organisms. Common amino acids might be expected to result in higher LCD frequencies, since encountering sufficient clusters of these amino acids is statistically more probable. A prototypical example is the *P. falciparum* proteome [37], which is remarkably N-rich and has a correspondingly high frequency of N-rich LCDs. As noted above, an abundance of particular types of LCDs may necessitate specific adaptations to cope with (or even leverage) those LCDs regardless of their underlying amino acid frequencies. Additionally, LCDs contribute to whole-proteome amino acid abundance, which could lead to underestimates of LCD enrichment since, in the absence of LCDs, the amino acid may be less abundant in the proteome. Nevertheless, examination of LCD frequencies in the context of whole-proteome amino acid frequencies provides a complementary view by indicating LCD classes that are more frequent than expected given the abundance of their constituent amino acids and the total number of proteins in the proteome.

To determine how often protein-level LCD frequencies exceed expectations based on background amino acid frequencies and proteome size, all proteins in every proteome evaluated were individually scrambled and re-evaluated for all LCD classes using LCD- Composer (see Methods). For each LCD class, LCD frequencies for the scrambled proteomes were then compared to corresponding LCD frequencies in the original proteome. Finally, for each LCD class, the percentage of organisms for which that LCD class was overrepresented relative to its scrambled frequency was calculated separately for each domain of life.

Relatively few LCD classes were significantly enriched across a high percentage of archaea or bacteria (Fig 4A,B; supplementary table available at [26]), which is due in part to the rarity of many LCD classes in these domains of life. Multiple LCD classes with A as the primary or secondary amino acid were significantly enriched in a reasonably large percentage of both archaea and bacteria. In addition to these classes, a number of D-rich LCD classes appear to be specifically enriched among archaea, whereas a number of bacteria exhibit enrichment of PA and AP LCDs. While overall percentages of viruses with LCDs are low for all LCD classes (Fig 4D), the highest values are achieved for the DG, DN, and PG classes. However, it is worth noting that a large percentage of archaeal and bacterial organisms exhibit at least some degree of LCD enrichment for a variety of LCD classes (Fig S6). In contrast, relatively few organisms exhibit LCD depletion (i.e., LCDs that are less common in the original proteome compared to the scrambled proteome; Fig S6), and statistically significant LCD depletion was extremely rare (Fig S7).

**Fig 4.**
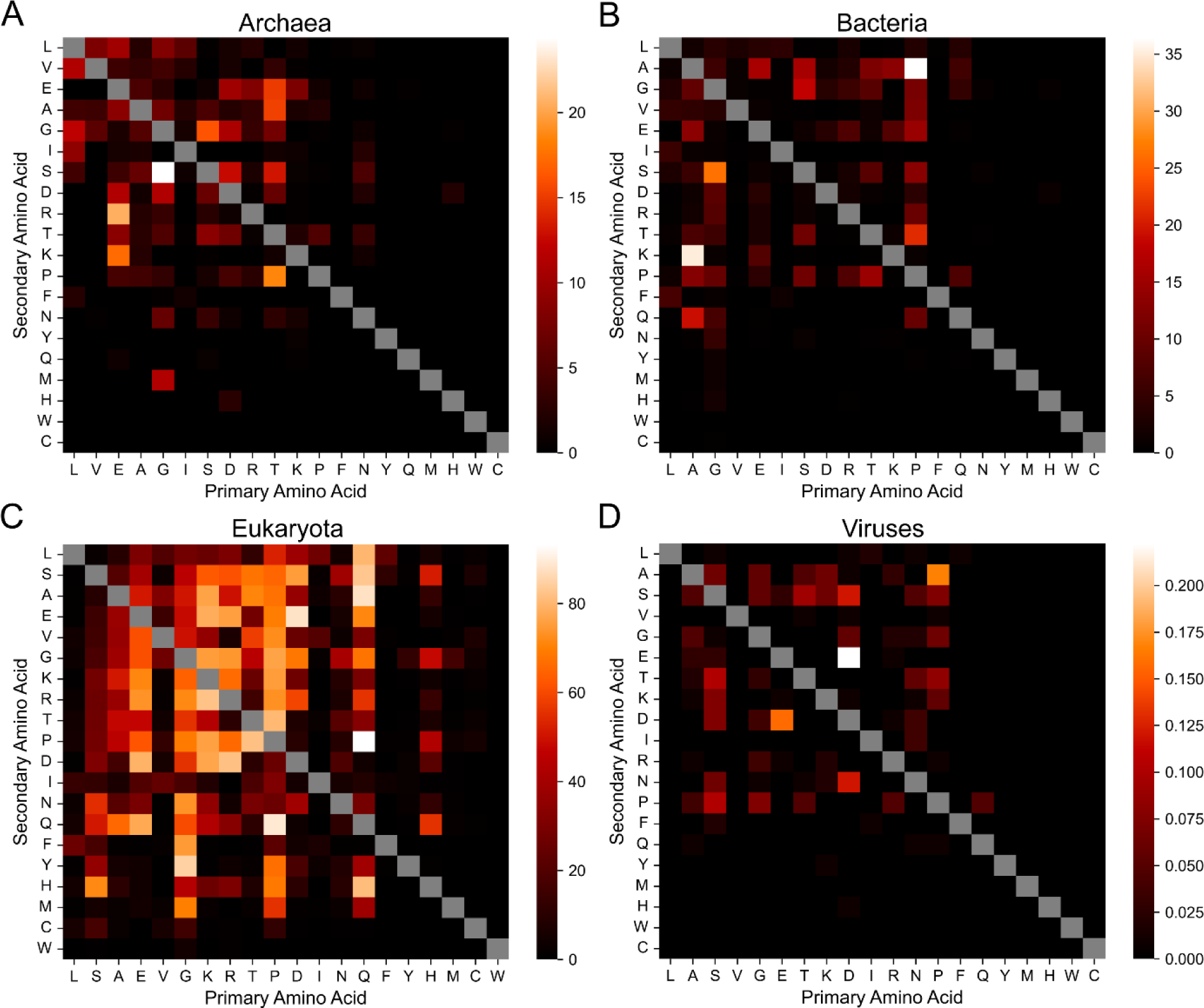
Percentage of organisms with significantly enriched LCDs after accounting for amino acid frequencies. The percentage of organisms with significantly enriched LCD-containing proteins (relative to a scrambled version of its proteome) is depicted for each LCD class in archaea (A), bacteria (B), eukaryota (C), and viruses (D). LCD frequencies in the original and scrambled proteomes were compared using Fisher’s exact test for each LCD class in each organism. Within each organism, *p*-values for all represented LCD classes (i.e., those with at least one LCD instance in the original or scrambled proteomes) were corrected using the Holm–Šidák correction method to account for multiple hypothesis testing. Significant enrichment is defined as *p*<0.05 after multiple-test correction.

However, it is worth noting that statistically significant depletion of LCDs (i.e. fewer LCDs in the original proteome compared to the scrambled proteome) was extremely rare across all LCD classes and domains of life [26].

In contrast to the other domains, eukaryotes exhibit both the highest percentages of organisms with significant LCD enrichment and the greatest variety of LCD classes achieving a high percentage (Fig 4C). High percentages tend to cluster among the 11 most common amino acids in eukaryotes, indicating that significant LCD enrichment typically occurs in spite of high background amino acid frequencies. However, amino acid frequency does not always determine LCD enrichment: L and V are ranked 1^st^ and 5^th^, respectively, in terms of amino acid abundance but their corresponding LCD classes are rarely enriched in eukaryotic organisms. Among the remaining 9 most abundant amino acids, specific secondary classes are enriched. For example, consistent with our previous observations in humans [38], SR LCDs are significantly enriched in many eukaryotes (90.5%), whereas SK LCDs – a related, positively charged class of LCDs – exhibit markedly lower enrichment (42.0%). This likely reflects the functional specificity of SR LCDs: while SR LCDs are strongly associated with RNA-binding and RNA-processing activities, SK LCDs are only weakly associated with these activities [38]. The reciprocal class, RS LCDs, are also enriched in a high percentage of organisms (86.5%), reflecting the flexible R/S composition criteria associated with SR-related proteins [38]. Similarly, GR and RG LCD classes are both significantly enriched across a high percentage of organisms (90.6% and 75.1%, respectively) while GK and KG LCD classes are significantly enriched in fewer organisms (24.0% and 33.2%, respectively), consistent with the functional importance and specificity of RGG domains [39]. However, enrichment of reciprocal classes is not always strong: QL LCDs are moderately enriched in many eukaryotes (32.7%), whereas LQ LCDs are notably less-frequently enriched (2.1%).

LCD classes composed of physicochemically similar amino acids are also not always similarly enriched across organisms and exhibit high context dependence. For example, RE and KE LCDs are frequently enriched in many eukaryotes (84.5% and 90.7%, respectively), whereas RD and KD LCDs are less-frequently enriched (58.6% and 11.3%, respectively). This difference may be due in part to the apparent compositional and structural preferences of charged single α-helices, where D may not be an effective substitute for E in this specific subclass of LCDs [40,41]. However, ED and DE LCDs are among the most enriched LCD classes (95.7% and 95.3%, respectively) and can play roles in regulating gene expression via nucleic acid mimicry [42,43], suggesting that these residues may be largely interchangeable, at least in some highly anionic LCDs. Conversely, KR and RK LCDs are less frequently enriched and exhibit a large difference in frequency of enrichment (37.4% and 12.5%, respectively), suggesting both that highly cationic LCDs are less common compared to highly anionic LCDs and that K and R are not interchangeable in this context.

Among the 9 least abundant amino acids in eukaryotes, LCD classes are significantly enriched across a high percentage of organisms less often. Only NS, HQ, QP, QA, QL, and QH classes reach significant enrichment in ∼15% or more of the eukaryotic organisms. Given the functional specificity of HQ and QH LCDs (Fig 3), the NS, QP, QA, and QL classes may also be selectively maintained and potentially utilized for specific functions across multiple organisms.

It is important to note that power in these experiments is limited both by LCD sample sizes (particularly for rare LCD classes, which have relatively large confidence intervals and high *p*-values) and by multiple-hypothesis test correction (since up to 400 tests are run for each organism). To examine whether the observed differences in LCD class enrichment were driven predominantly by underlying differences in statistical power, the typical degree of enrichment for each LCD class was estimated as the median natural logarithm of the odds ratio (lnOR) across organisms (see Methods). Importantly, the median lnORs for the LCD classes composed of common amino acids are also higher compared to the median lnORs for LCD classes composed of less-common amino acids (Fig S8; supplementary data available at [26]), suggesting that differences in the percentage of organisms with significant enrichment across LCD classes is not primarily driven by differences in statistical power (though it is clearly still a contributor). This method can also be applied to individual organisms: the malarial parasite proteome – a prototypical proteome with highly skewed amino acid frequencies – exhibits significant enrichment across a variety of N-rich LCD categories despite the unusually high N frequency within its proteome (Fig 5A,B). Similarly, E and D are the 5^th^ and 6^th^ most common amino acids, respectively, yet a variety of E-rich and D-rich LCD classes are among the most strongly enriched in the malarial proteome – classes that are understudied in *P. falciparum* compared to N-rich LCDs. For example, the native malarial proteome contains 228 proteins with at least one DE LCD (4.2% of the proteome), but none are retained after scrambling. In humans, with the exception of L and V, the LCD categories corresponding to the most frequent amino acids exhibit significant enrichment across a variety of secondary LCD classes (Fig 5C,D). Additionally, although whole proteome frequencies for H and C are relatively low in humans and power is generally lower for these LCD classes, multiple LCD classes are significantly enriched in the proteome. Thus, LCD instances are not solely attributable to amino acid frequencies, even in organisms with highly biased background amino acid frequencies.

**Fig 5.**
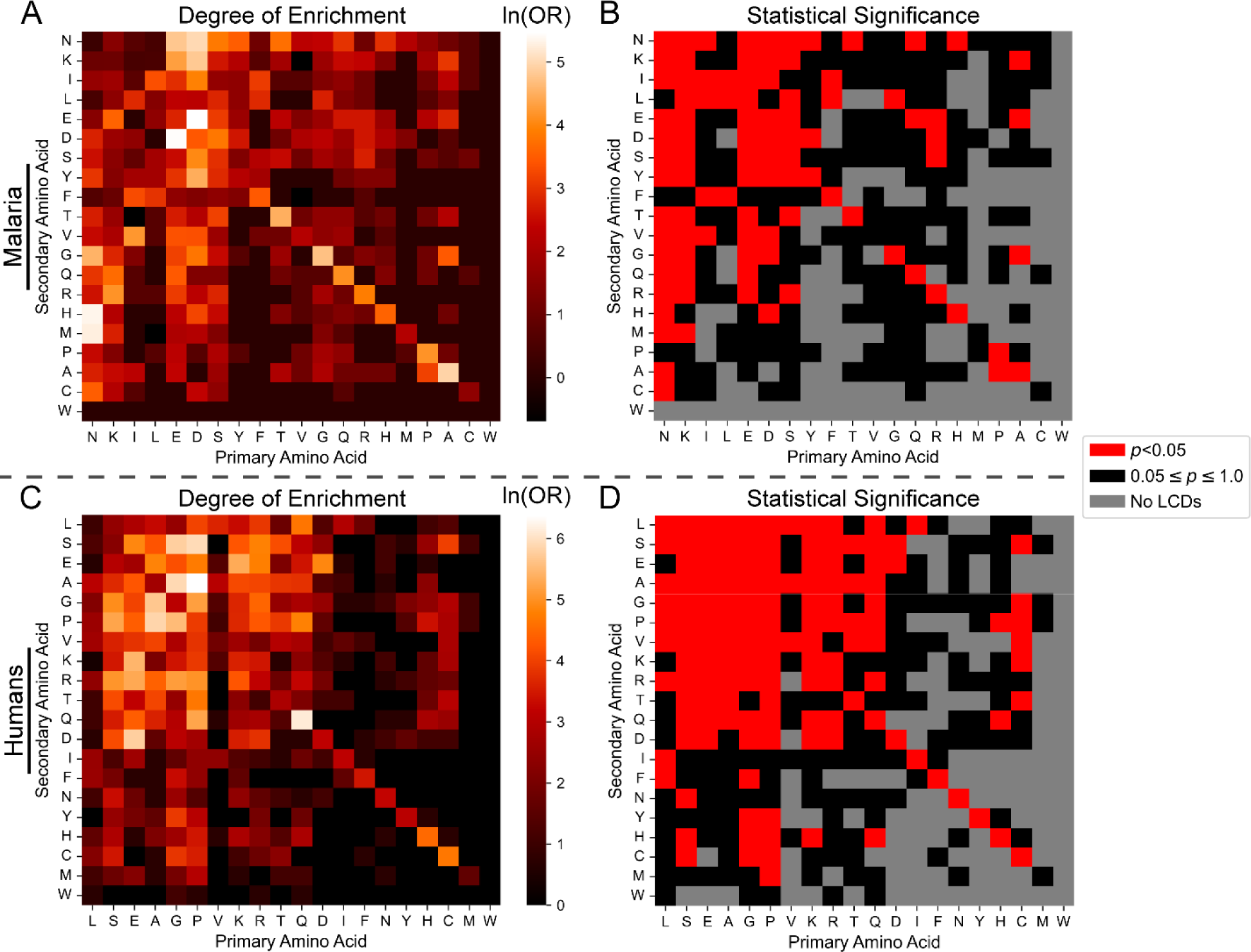
Statistical LCD enrichment by LCD class in the malarial and human proteomes. (A) Heatmap depicting the degrees of LCD enrichment (expressed as the lnOR) for each LCD class among the *P. falciparum* proteome (UniProt ID: UP000001450_36329). For LCD classes in which the number of LCDs in either the original or scrambled proteomes were 0, a value of 1 was added to all cells in the contingency table to calculate a biased lnOR (see Methods). (B) Binary classification for LCD categories for which enrichment was statistically significant (red squares) or statistically non-significant (black squares) after multiple-test correction. Grey squares indicate LCD categories that were excluded from statistical analysis since no LCDs were found in both the original and scrambled proteome. (C) Degrees of LCD enrichment in the human proteome. (D) Statistical significance for LCD enrichment in the human proteome. For all panels, the diagonals represent corresponding values for each primary LCD class. The data underlying these heatmaps can be found in the supplementary data available at [26].

In summary, domains of life exhibit both shared and unique enrichment for specific LCD classes. Eukaryotes exhibit the greatest breadth and most consistent enrichment for a variety of LCD classes, with many enriched classes corresponding to those with known (e.g., SR/RS, GR/RG, DE/ED) or new (e.g., HQ) functional significance. Importantly, LCD classes almost universally exhibit degrees of enrichment that are greater than or equal to those expected based on whole-proteome amino acid frequencies, suggesting that they are often not simply byproducts of amino acid frequencies.

### Per-residue-level LCD Class Frequencies: Select Organisms Achieve Highest Per-Residue Occupancy for Multiple LCD Classes

The measure of total proteins with LCDs for each LCD class in each organism identifies proteomes with an unusually large number of LCD-containing proteins and aids in determining putative LCD class functions (which relies on non-redundant sets of proteins). While this statistic provides an enlightening and useful view of extreme LCD content, it does not account for multiple LCDs within a single protein or the lengths of LCDs within each protein, and organisms with large proteomes are often selectively enriched among the top-ranking organisms. A complementary measure of LCD content is the per-residue occupancy of LCDs, defined as the percentage of all residues in each proteome that are located within LCDs for each class. This statistic offers a slightly different view of whole-proteome LCD content by effectively estimating the percentage of proteome space devoted to each LCD class within each organism.

We first identified organisms with the most extreme per-residue occupancy for each LCD class. For many primary LCD classes, per-residue occupancy often reached single-digit or even double-digit percentages for the top three organisms from all domains of life and were sometimes orders of magnitude higher than the mean per-residue occupancy across all proteomes within the corresponding domain of life (Fig 6, Table S9). Some LCD classes have high per-residue occupancy values among the top scoring organisms from all four domains of life, while certain domains of life exhibit uniquely high maximum per-residue occupancies for specific LCD classes. For example, the top-ranking organisms achieve relatively high per- residue occupancies for A-rich LCDs across all domains. The highest per-residue occupancy for G-rich LCDs is similar for bacteria and eukaryotes but low among archaea. Eukaryotes exhibit uniquely high per-residue occupancies for N-rich, P-rich, Q-rich, and S-rich LCDs compared to archaea and bacteria. For most of these classes, higher per-residue occupancy is achieved despite larger proteomes than is typical for archaea and bacteria (3203 proteins, 25399 proteins, 7635 proteins, and 11530 proteins for the top-ranking proteome in the N-rich, P-rich, Q-rich, and S-rich categories, respectively).

**Fig 6.**
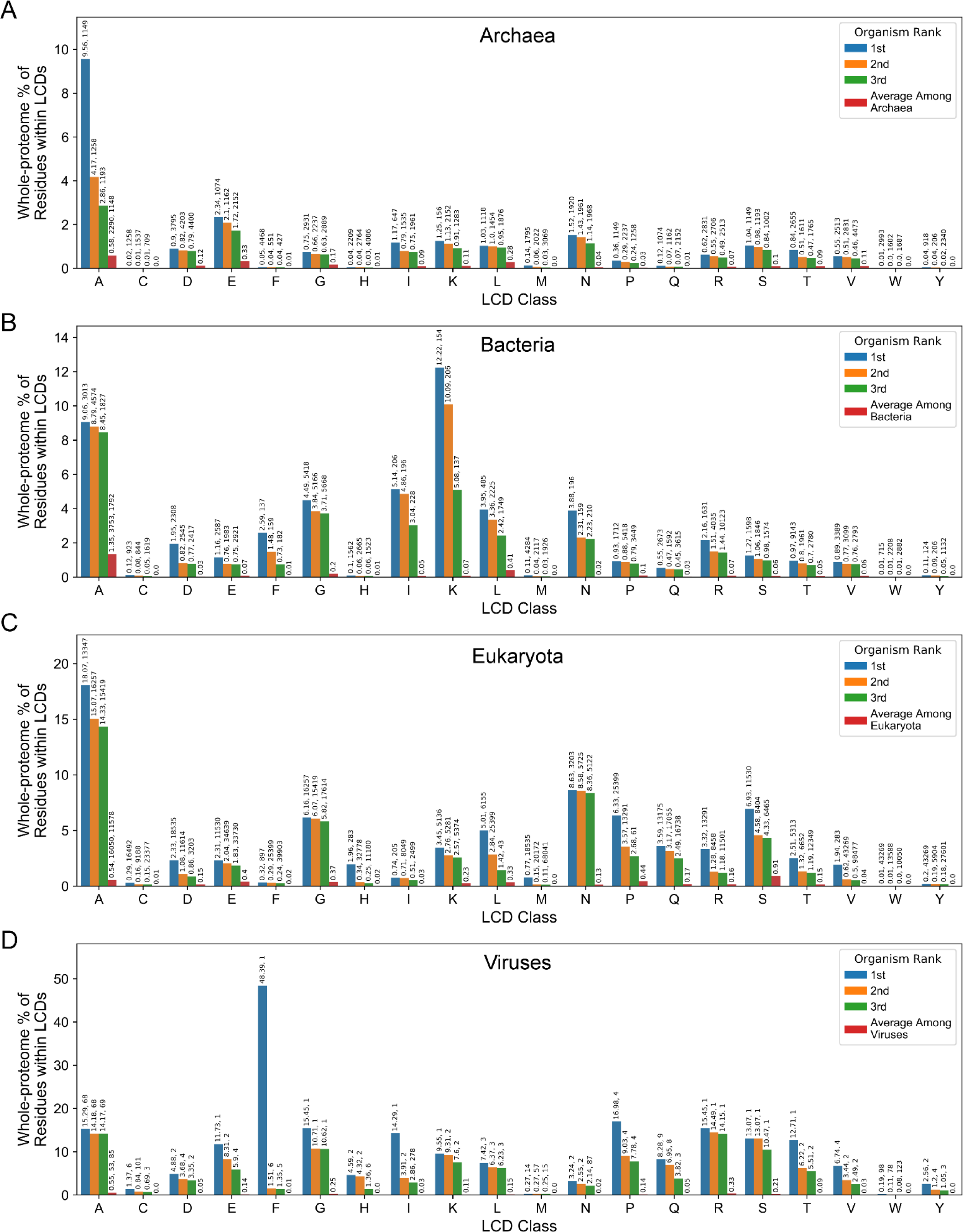
Per-residue occupancy for the top-ranking organisms from each domain of life for the primary LCD classes. Per-residue occupancy was calculated separately for each organism as the percentage of total residues in the proteome that were occupied by LCDs from each primary LCD class for archaea (A), bacteria (B), eukaryota (C), and viruses (D). Values above each bar represent the per-residue occupancy value (as a percentage), followed by the total number of proteins in the corresponding organism. For the average among each domain of life (red bars), the mean number of proteins pertyts organism and the standard deviation in the number of proteins per organism is expressed above the bar for the “A” LCD class only since these values are independent of LCD class.

Viruses tended to exhibit some of the highest per-residue occupancy values across many LCD classes, but this often occurs in exceptionally small proteomes. For example, the two extremely high values – 48.4% occupancy for F-rich LCDs and 27.9% occupancy for I-rich LCDs – occur in the Cotton leaf curl Multan betasatellite and Bhendi yellow vein mosaic betasatellite, respectively. Both betasatellites depend on helper viruses for their replication and encode a single protein, making their per-residue occupancy values unusually large outliers even among this set of maximum values. Similarly, some of the highest per-residue occupancy values among other domains (especially bacteria), correspond to extremely small – and potentially incomplete – proteomes. Furthermore, although the top-ranking eukaryotic proteome for F-rich LCDs is reasonably large (>50k proteins, corresponding to the pharaoh cuttlefish, *Sepia pharaonis*, its per-residue occupancy of F-rich LCDs is so high (18.5% of the proteome) that we consider it unlikely to be a legitimate proteome and we excluded it from Fig 6 and all subsequent analyses. Therefore, while the study of organisms with the highest LCD content is informative and interesting, we emphasize caution in interpreting values for small proteomes and extremely unusual outliers.

We next calculated the per-residue occupancy of each secondary LCD class for each organism and focused on those with the highest values for each LCD class. Differences between specific groups of LCD classes become more prominent when per-residue occupancy is considered (Fig 7 and Table S9). For instance, only select NX classes (i.e., those with N as the primary residue) – including the NI, NS, ND, NT, and NK classes – have relatively high maximum per-residue occupancies in eukaryotes. High per-residue occupancies for NX classes are limited to the NI, NS, NT, and NK classes in archaea, and high values in bacteria are further limited to the NI and NK classes only. As another example, high per-residue occupancies are achieved for PL, PS, PA, PG, and PR classes (all being types of PX classes), whereas PX classes do not reach high per-residue occupancies in archaea or bacteria, and the highest PX values are observed for different PX classes.

**Fig 7.**
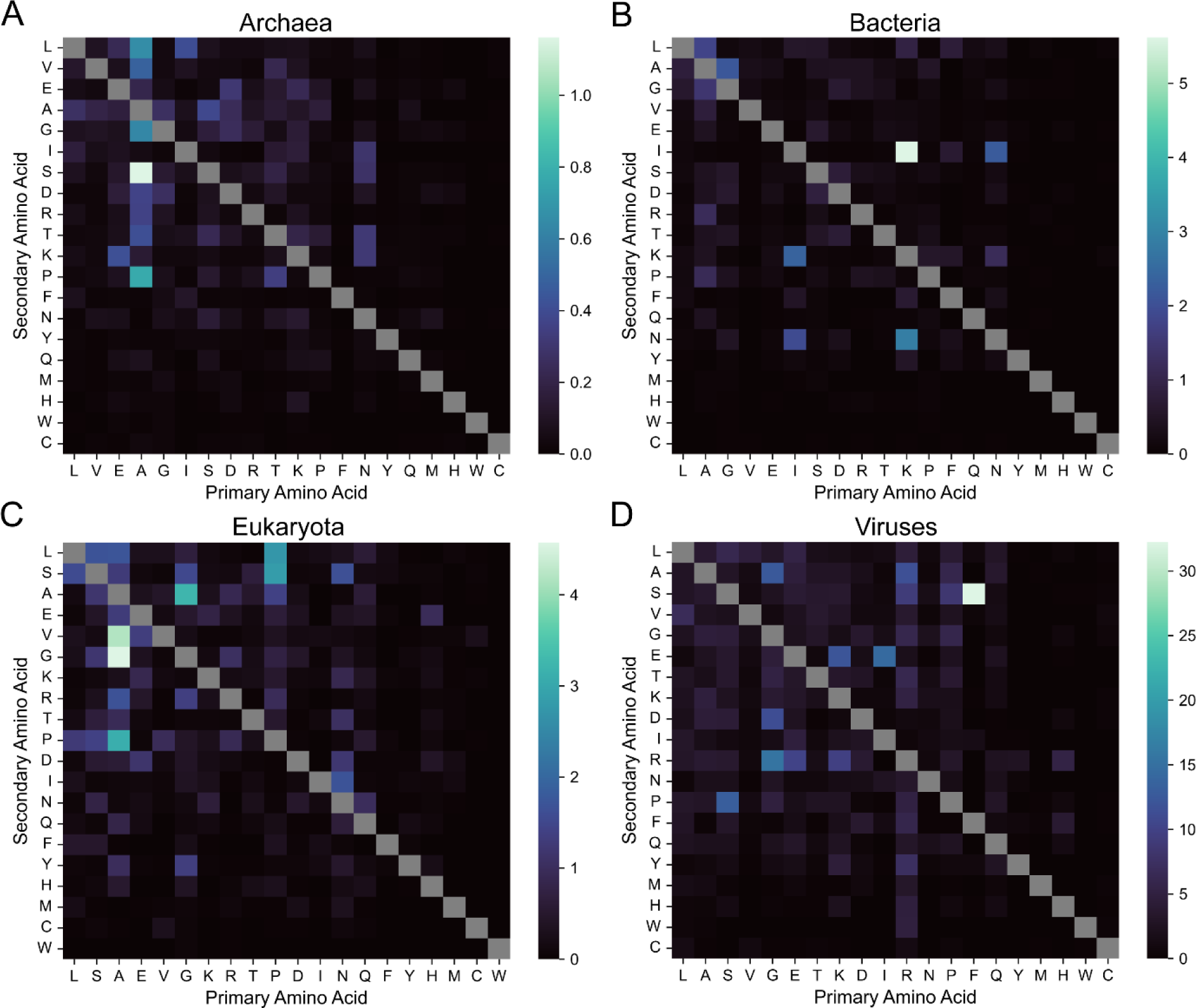
Maximum per-residue occupancy for each LCD class by domain of life. Per-residue occupancy was calculated for each LCD class and each organism. Maximum per-residue occupancy is depicted separately for each LCD class in archaea (A), bacteria (B), eukaryotes (C), and viruses (D).

Certain organisms have the absolute highest per residue occupancy for multiple LCD classes, suggesting that these proteomes are extremely unusual. For example, in eukaryotes, ∼18% of the 400 LCD classes are attributed to only five organisms (Fig 8A). A closer examination of these organisms reveals organism-specific clusters of LCD classes with maximum per-residue occupancy values (Fig 8B). For example, *S. microadriaticum* – a dinoflagellate important in coral reef ecosystems [44,45] – contributed the maximum per-residue occupancy for 19 separate LCD classes (the most from any single organism), and these classes tended to correspond predominantly to LCDs with hydrophobic or aromatic amino acids as the primary residue. The whiteleg shrimp, *Penaeus vannamei*, had the highest per-residue occupancy for multiple LCD classes with L, F, P, or S as the primary amino acids. A species of seaweed (purple laver; *Porphyra umbilicalis*) exhibits per-residue occupancies that are specifically high for LCD classes with R as the primary or secondary amino acid, along with a small number of P-rich or G-rich LCDs. The intestinal fluke, *Echinostoma caproni*, achieves the highest per-residue occupancy for multiple LCD classes with M or D as the primary amino acid. Finally, a species of malaria (*Plasmodium falciparum RAJ116*) exhibits the expected highest per residue LCD occupancy for multiple classes with N as the primary residue, along with a few additional classes containing D or Y as the primary residue. Similar LCD-class specificity is also observed in the top ranking archaeal, bacterial, and even viral organisms (Fig S9).

**Fig 8.**
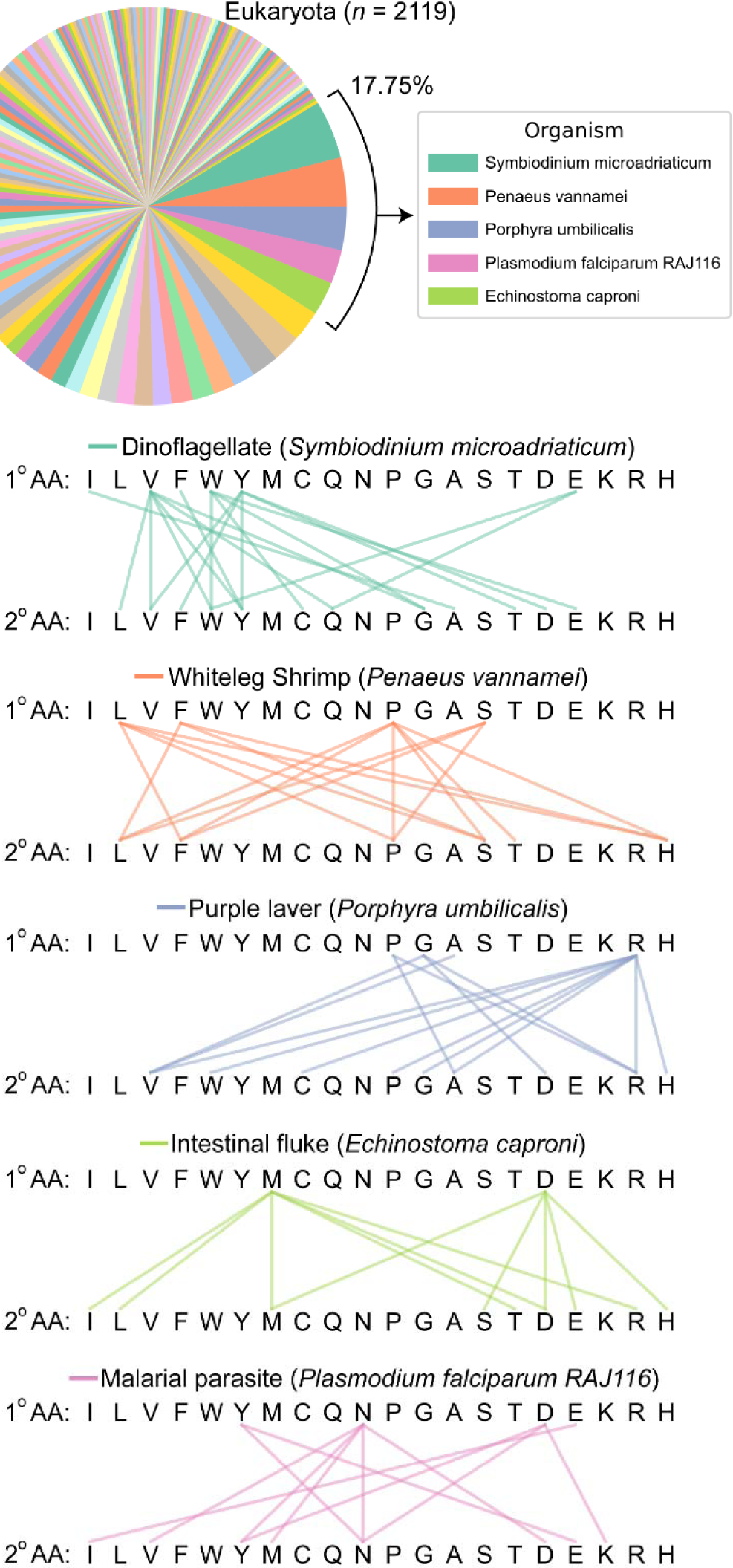
Number of LCD classes assigned to each eukaryotic organism contributing a maximum per-residue occupancy for at least one LCD class. (A) Pie chart indicating the assignment of LCD classes (400 total) to the eukaryotic organism achieving the highest per-residue LCD occupancy. Each wedge represents a single organism associated with the overall highest per-residue occupancy observed among eukaryotes. Wedge size indicates the number of LCD classes for which the single organism corresponding to that wedge achieved the highest per-residue occupancy. The top five eukaryotic organisms are indicated in the legend. Out of necessity, the color palette was repeated in the pie chart, though each color cycle represents a different set of organisms. (B) Linkage maps indicate the types of LCD classes for which the organism contributed the maximum per-residue occupancy value for eukaryotes. The first row of amino acids in each linkage map indicates the primary amino acid comprising the LCD class, and lines connected to the second row of amino acids indicate the secondary amino acid comprising the LCD class. Lines connecting identical amino acids (e.g., W connected to W) indicate that the organism contributed the maximum per-residue occupancy value for the primary LCD class as a whole (e.g., the W-rich primary LCD class). LCD classes without connecting lines are those for which the organism did not contribute the maximum per-residue occupancy value. Similar analysis for archaea, bacteria, and viruses can be found in Fig S9. Although it achieved high per-residue occupancies for multiple LCD classes, the *Spodoptera litura* proteome was manually identified as an exceptionally incomplete reference proteome and excluded from analyses.

Collectively, these observations highlight the extreme percentage of proteome space devoted to each class of LCD in specific organisms, uncover organism-specific preferences for certain LCDs, and identify organisms with exceptionally unusual proteomes.

### LCD-Centric Comparison of Whole Proteomes via Analysis of LCD Class Distributions and Per-Residue LCD Occupancy

We have shown previously that primary LCD frequencies vary across organisms, resulting in unique LCD frequency profiles [4]. By extension, we expected secondary LCD frequencies to also vary across organisms. In order to perform a direct cross-organism comparison of the distributions of secondary LCDs within each primary LCD class, all instances of secondary LCDs were collected for each organism. For each secondary LCD class, the “share” of secondary LCDs was calculated as the number of LCDs in that class divided by the total number of secondary LCDs with the same primary amino acid, expressed as a percentage (e.g., the share of HQ LCDs is the percentage of HQ LCDs out of all HX LCDs).

Differences are apparent both when comparing two individual organisms and when comparing a single organism to all organisms in the same domain of life (Fig 9). For example, human A-rich secondary LCDs exhibit expanded shares devoted to AP and AG classes compared eukaryotes in general (Fig 9A,B), with a corresponding contraction of other A-rich classes in humans relative to eukaryotes. Additional differences are observed among other classes: for example, VL, HP, and DE classes are all expanded within their respective primary LCD categories in humans relative to eukaryotes, necessitating compensatory contraction of other secondary classes in the same primary class. Similar types of comparisons can be made between the LCD profiles for individual organisms. For example, in humans P very frequently serves as the secondary amino acid across a variety of primary LCD classes, whereas P is substantially less common as the secondary amino acid in yeast (Fig 9B,C). Conversely, N frequently serves as the secondary amino acid in yeast but not in humans. While other differences are also readily apparent, these representative examples highlight the variation in LCD content profiles across organisms.

**Fig 9.**
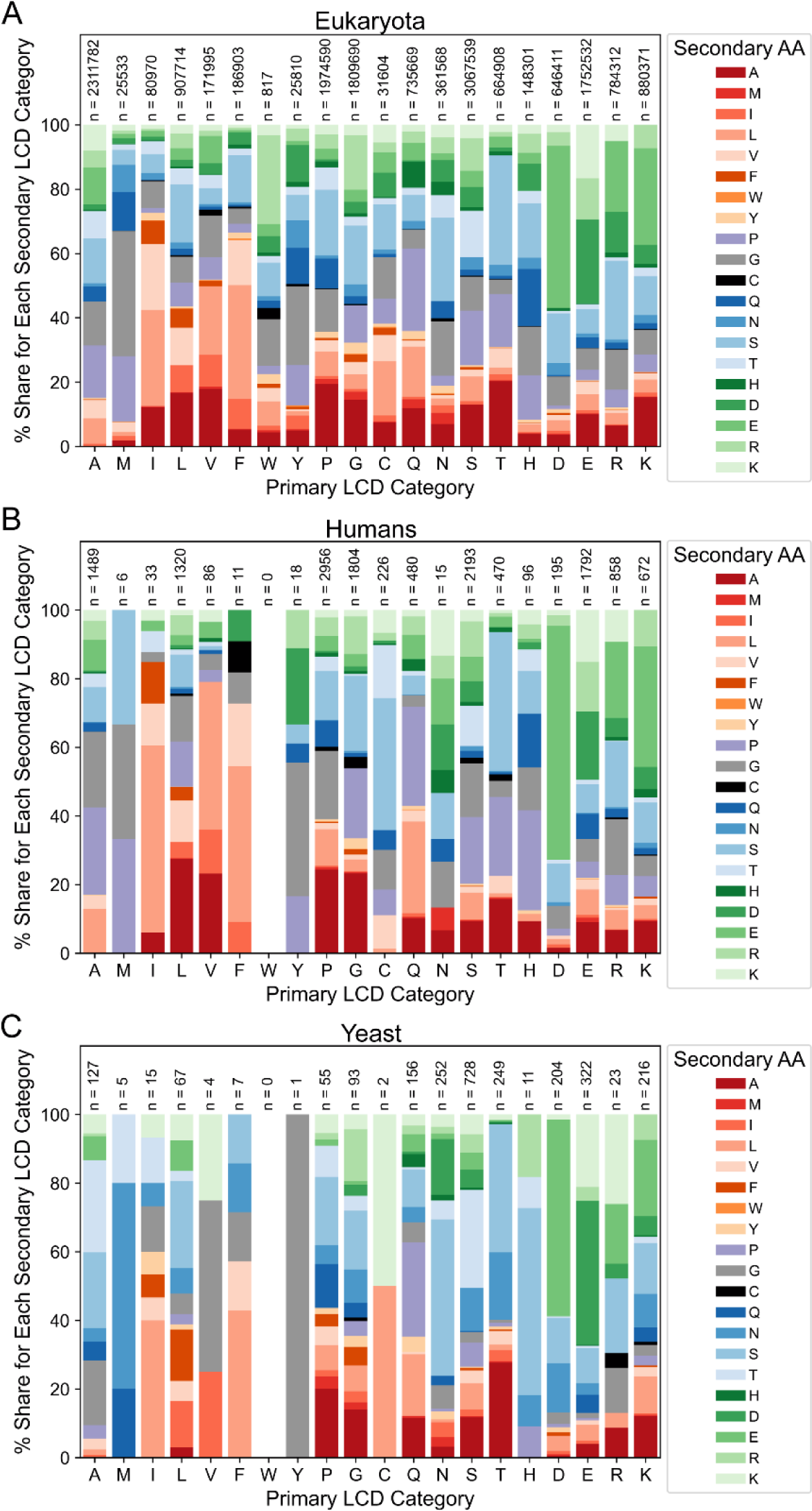
Comparison of secondary LCD distributions among eukaryotes. (A) Distributions of secondary LCDs within primary LCD categories for all secondary LCDs identified in eukaryotic organisms. (B) Distributions of human secondary LCDs among primary LCD categories. (C) Distributions of yeast secondary LCDs among primary LCD categories. For all panels, secondary LCDs were grouped into a primary LCD category based on the predominant amino acid used in the LCD search. Then, within each primary LCD category, the percentage of total LCDs in that category was calculated for all possible secondary LCD categories and depicted as a stacked bar plot. For all secondary amino acid classes, the primary amino acid is represented on the *x*-axis, the bar color specifies the secondary amino acid, and the size of the bar indicates the percentage. Secondary amino acids are loosely grouped and colored according to physicochemical properties and appear in the same order (from bottom to top) as in the figure legends. Total LCD sample sizes are indicated above each bar.

Quantification of LCD content as per-residue occupancy also facilitates equitable cross-proteome comparisons since it normalizes for proteome size and accounts for differences in LCD lengths. LCD content for each proteome can be represented by a 400-attribute array of per-residue occupancy values corresponding to the 20 primary LCD classes and 380 secondary LCD classes. Data in this work as well as prior work [4,5,46–50] suggests that organisms exhibit characteristic “signatures” of LCD content. With this classification scheme, LCD-content signatures could be used to compare organisms. As a representative example, the LCD-content signature – expressed as the percentile rank for every LCD class compared to all eukaryotes – of the human proteome was compared to the LCD-content signature of *Saccharomyces cerevisiae* (budding yeast). Humans exhibit high percentile ranks for the P, R, G, L, C, and E primary LCD classes, indicating that these classes occupy a larger percentage of the human proteome compared to most eukaryotes (Fig 10A). In striking contrast, yeast exhibits high percentile ranks for the Q, T, K, S, I, F, M, D, and N primary LCD classes. Other LCD classes exhibit little or no difference in per-residue occupancy, likely because many of them are not common in both yeast and humans.

**Fig 10.**
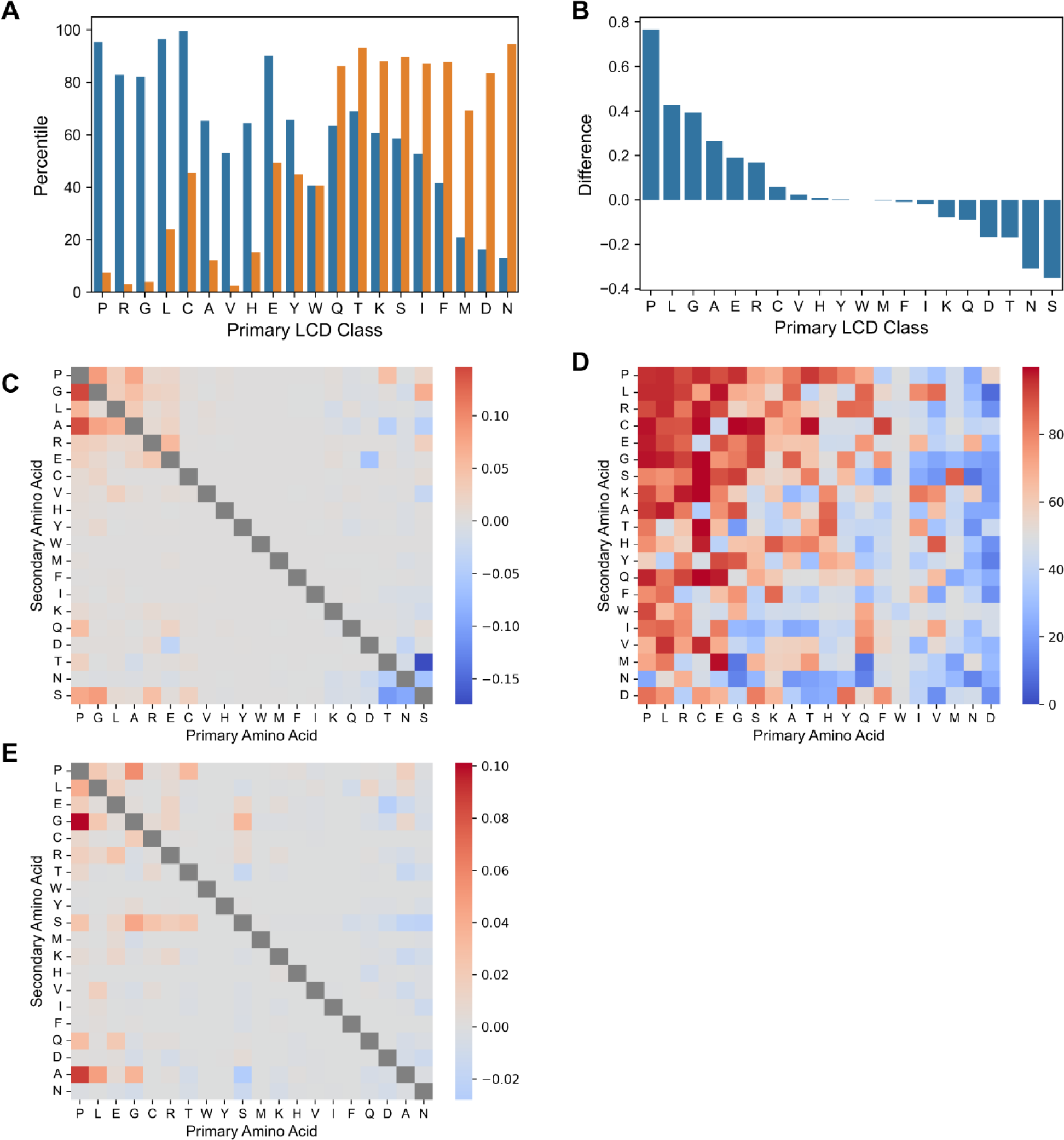
Comparison of organisms based on whole-proteome per-residue occupancy of LCDs. (A) Percentile rank for each of the primary LCD classes in humans (blue) and yeast (orange) relative to all eukaryotes. The primary LCD classes (x-axis) are sorted by difference in percentile from largest to smallest. (B) Raw difference in per-residue occupancy for each of the primary LCD classes in humans compared to yeast. (C) Raw difference in per-residue occupancy between humans and yeast for each of the secondary LCD classes. (D) Percentile rank for each of the secondary LCD classes in humans relative to all eukaryotes. (E) Raw difference in per-residue occupancy between humans and the corresponding average value among eukaryotes for each of the secondary LCD classes.

Raw per-residue occupancy values can also be directly compared, illuminating the actual magnitude of differences in LCD content between organisms. For example, by percentile rank, yeast and humans differ substantially with respect to N-rich LCD content but only modestly with respect to S-rich LCD content (Fig 10B). However, by raw difference in per-residue occupancy, the difference between humans and yeast is more extreme with respect to S-rich LCDs than N-rich LCDs (Fig 10A), likely reflecting, at least in part, the abundance of S-rich LCDs in yeast cell wall proteins [46]. Indeed, among the secondary LCD classes, the most negative raw difference in per-residue occupancy (indicating higher LCD content in yeast relative to humans) occurs specifically for the ST and TS classes common among yeast cell wall proteins, whereas humans actually exhibit a higher per-residue occupancy for the SG, SR, and SP classes (Fig 10C).

In addition to pairwise comparisons between organisms, individual organisms can be compared to the average per-residue occupancy among all other organisms of the corresponding domain of life. Again, both the percentile rank and the raw difference in per-residue occupancy relative to the mean offer complementary comparisons. Humans achieve a very high percentile rank for nearly all P-rich secondary LCD classes (Fig 10D), indicating that these classes have unusually high per-residue occupancy in humans relative to other eukaryotes. However, PG LCDs (and, to a lesser extent, PA LCDs) clearly have the highest raw difference in per-residue occupancy relative to the mean (Fig 10E), highlighting the LCDs classes for which humans differ from other eukaryotes by the greatest magnitude.

## Discussion

Our study was founded upon four related guiding questions: how much does LCD content vary across organisms, where is each type of LCD found, how common is each type of LCD, and what are the functions associated with LCDs (particularly rare or niche types of LCDs)? We find that even the most unusual categories of LCDs are present in nature, sometimes even at extremely high levels in particular organisms. More broadly, organisms sometimes differ dramatically with respect to their overall LCD content profiles. In some organisms, biased genome compositions contribute substantially to LCD frequencies. However, we find that LCDs are often significantly enriched despite these biases. Furthermore, organisms must have ways to leverage (or at least tolerate) such unusual features of their proteomes regardless of their origins.

By definition, our inquiry into rare and unusual LCDs is, to some degree, a foray into understudied corners of biology. One of the limitations of this study is the paucity of information and the difficulty in experimentally characterizing non-model organisms. As a result, we have relied heavily on computational tools, predictions, and the limited information currently available. Although the existing proteome resources are extremely well-maintained, some of the proteomes in our dataset await detailed experimental validation, including direct detection of all proteins *in vivo*. Therefore, while the sequences may indicate the presence of LCDs, their existence and abundance in the corresponding organism may not be fully known. Our intention is for this study to serve as a launch-point for further investigation.

LCDs of some types were found at reasonably high levels in well-studied organisms, enabling their examination at greater depths. For example, HQ LCDs were present in multiple organisms (including two model organisms, *D. melanogaster* and *D. discoideum*, with high HQ LCD content) and were associated with strongly overlapping sets of transcription and RNA-related functions. Q/H-rich LCDs can serve as pH sensors in multiple organisms through alterations in protein structure and activity [15,16], even when H levels are not particularly abundant within the LCD [51]. It is tempting to speculate that HQ LCDs may be widely used, pH-responsive LCDs commonly associated with transcription and gene expression, as observed recently for the yeast transcription factor, Snf5 [51], and the mRNA-binding and translation-regulating Orb2 protein in *Drosophila* [15,16].

Keratin proteins emerged as an exemplar family of eukaryote-specific proteins with distinct subcategories of LCDs. Multiple C-rich secondary LCD classes were represented in a relatively large number of proteins in a variety of well-studied eukaryotic organisms. Given their purpose as structural proteins in hair, skin, nails, claws, and other related features, these LCDs were sensibly identified as eukaryote-specific LCDs. However, this also suggests that these types of LCDs are not frequently utilized for other functions in non-eukaryotic organisms.

Additionally, LCD-Composer provided the specificity to distinguish between proteins of the same keratin family with distinct classes of LCDs and LCD profiles, while also detecting overlap between LCD classes among keratins. The high number of proteins with these LCDs in well-studied organisms, along with the specialization of certain categories of LCDs among the keratin family, made keratins a useful model for eukaryote-specific LCDs and even revealed organism-specific utilization of distinct balances of LCDs among keratins in each organism.

However, not all C-rich LCDs were associated with keratin-related functions: multiple classes of C-rich LCDs were associated with different functions in different organisms (notably, CK LCDs with functions in metallothioneins).

Functional inferences for LCD classes were made primarily based on GO-term analyses.

While this method is widely implemented and is clearly useful (as exemplified by the C-rich classes of LCDs), it is not without limitations. First, enriched GO terms are simply statistical associations between groups of proteins and particular functional annotations. Not all proteins of a group will have the specified function, so caution should be exercised when extrapolating these associations to the same LCD classes in different organisms. Second, LCD function can differ between organisms and contexts. As an interesting example, a mammalian LCD normally involved in alternative splicing, protein-protein interaction, and maintaining protein solubility [52] was bioengineered to act as extremely strong and environmentally resilient extracellular adhesive protein in mussels [53]: a function that is only tangentially related to its native functions. Third, biases inherent in the gene ontologies may limit the functional associations that can be observed or that reach statistical significance. Finally, it is possible that there are length-dependent relationships between LCD occurrence, the corresponding sets of LCD-containing proteins, and their associated functions. LCDs may be more likely to occur in longer proteins, which may, in-turn, be related to specific functions [4,25]. However, LCDs may also occur in these proteins for functional reasons, making it difficult to disentangle length-to-function relationships. We observe clear specificity in the functions associated with distinct classes of LCDs, suggesting that length-independent relationships between LCDs and associated protein functions are still easily identifiable.

In summary, our comprehensive survey of LCDs in the reference proteomes of all known organisms provides a powerful resource and classification system for LCDs. These LCDs may serve context-specific, adaptive functions in their native organisms, which may be potentiated by environmental conditions and/or concomitant proteome adaptations. Some organisms exhibit extremely unusual proteomes with hundreds of proteins containing rare types of LCDs, or with the highest LCD content for multiple classes of LCDs. In essence, a realized (i.e., expressed) proteome represents a major subset of the intracellular and extracellular ecosystem of biomolecules in a given organism. Considering the unusual nature of some LCD content profiles, these observations stretch our understanding of biomolecular ecosystems compatible with life.

## Methods

### Data Acquisition and Processing

UniProt reference proteomes for all available organisms were downloaded from the UniProt FTP site (https://ftp.uniprot.org/pub/databases/uniprot/) on 8/22/22. The gene ontology was downloaded from the Gene Ontology Consortium website (http://geneontology.org/docs/download-ontology/). Pfam annotations were acquired from the Pfam FTP site (http://ftp.ebi.ac.uk/pub/databases/Pfam/). Gene annotation files for GO-term analyses were downloaded from the European Bioinformatics Institute website (ftp://ftp.ebi.ac.uk/pub/databases/GO/goa/). GO-term analyses were performed using GOATOOLS [54] with default parameters using the same proteomes analyzed in the initial LCD searches.

### LCD Identification and Classification

Primary LCDs were identified using LCD-Composer with default parameters [4] for each reference proteome. Briefly, a 20-amino acid sliding window was used to scan each protein sequence. Windows passing the default minimum 40% composition corresponding to a single amino acid and a minimum linear dispersion of 0.5 were classified as LCDs. All overlapping windows passing these criteria were merged to generate contiguous LCDs. LCD searches were performed separately for each of the 20 canonical amino acids.

Secondary LCDs were identified using a separate LCD-Composer search for all possible combinations of two amino acids except those where the first and second amino acid are the same (380 total combinations). For each search, the minimum composition criteria were ≥40% and ≥20% for the first and second amino acid, respectively, comprising the secondary LCD class. For example, HQ LCDs were defined as those with a minimum of 40% H and a minimum of 20% Q. Searches were performed with a 20-amino acid window size and a minimum linear dispersion of 0.5 applied separately for each of the amino acids in the search.

A database of all primary and secondary LCDs identified as part of this study is available in a publicly accessible repository [26].

### Pfam Domain Mapping and Quantification

Pfam domain annotations were mapped to LCD-containing proteins for each class across all organisms. Pfam domains were then assigned to “clans” to represent the broadest possible categories linking the more specific domain annotations. For each organism, the maximum number of LCD-containing proteins associated with a single Pfam clan was calculated for each LCD class, only considering LCD classes for which at least one protein was assigned a Pfam annotation. These values were then averaged across organisms within each domain of life.

### Proteome Scrambling and LCD Frequency Estimation

Each proteome in the dataset was scrambled once using the Fisher-Yates shuffle method. To best represent constraints of true proteomes, the starting residue of each protein (typically a methionine) was extracted prior to scrambling. The scrambled proteome was then segmented into proteins of sizes equal to those found in the corresponding non-scrambled proteome, and the initiator residue was added back to the start of each scrambled protein. LCD- Composer was run on each scrambled proteome to identify both primary LCDs and secondary LCDs for every LCD class, exactly as performed for the original proteomes. The number of proteins containing an LCD was tallied for each scrambled proteome and LCD class, then compared to the corresponding value from the original proteomes using Fisher’s exact test.

Within each organism, *p*-values were adjusted using the Holm–Šidák correction method to account for multiple-hypothesis testing. The natural logarithm of the odds ratio (lnOR) and its corresponding 95% confidence interval were also calculated. When evaluating individual organisms, *P*-values corresponding to LCD classes with 0 instances in both the original and scrambled proteomes for a given organism were not calculable and were therefore excluded from multiple-test correction. However, when calculating the percentage of organisms with significant enrichment of LCDs from each LCD class, these classes were retained and considered not statistically significant (Fig 4). The median lnOR was used to estimate the typical degree of enrichment for each LCD class across organisms within each domain of life (Fig S8).

For LCD classes in which the number of proteins containing an LCD in either the original or scrambled proteomes was 0, a value of 1 was added to each cell in the contingency table to provide a biased lnOR estimate. Importantly, when a biased estimate was required, it was almost universally due to 0 LCD occurrences in the scrambled proteome: therefore, in the vast majority of cases, this procedure results in conservatively biased lnORs that underestimate the degree of enrichment observed in the original proteomes.

## Supplementary Data

**Fig S1.**
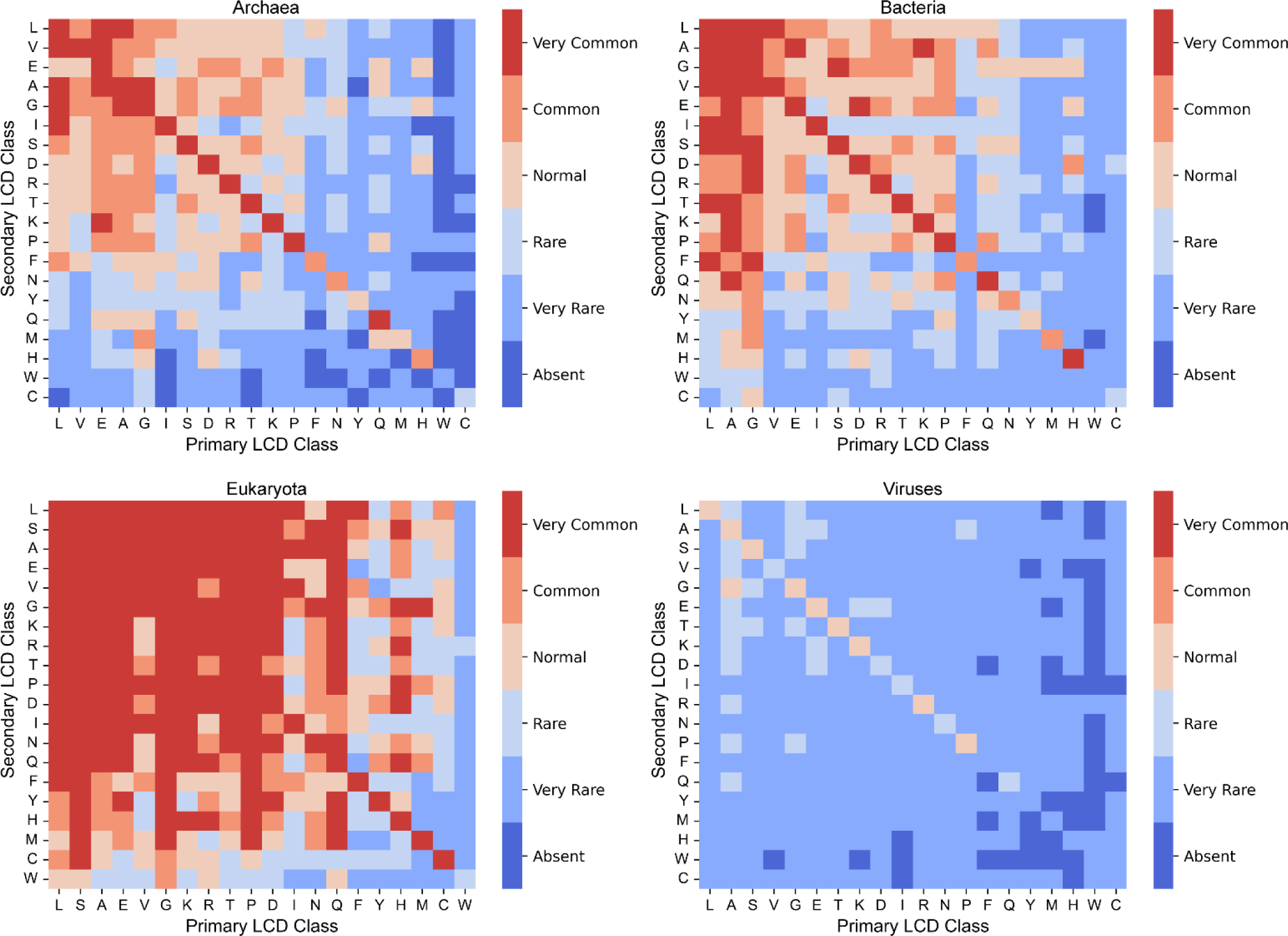
LCD frequency classification across the domains of life. Each type of LCD was classified as absent (*x*=0%), very rare (0%<*x*<5%), rare (5%≤*x*<20%), normal (20%≤*x*<50%), common (50%≤*x*<75%) or very common (*x*≥75%) separately for each domain of life, where *x* represents the percentage of organisms containing at least one instance of the LCD class. Squares on the diagonal represent the primary LCD classes, whereas off-diagonal squares represent secondary LCD classes. For each domain of life, amino acids on the axes are ordered from most-common to least-common based on the mean rank of whole-proteome frequency for each amino acid across all proteomes for that domain.

**Fig S2.**
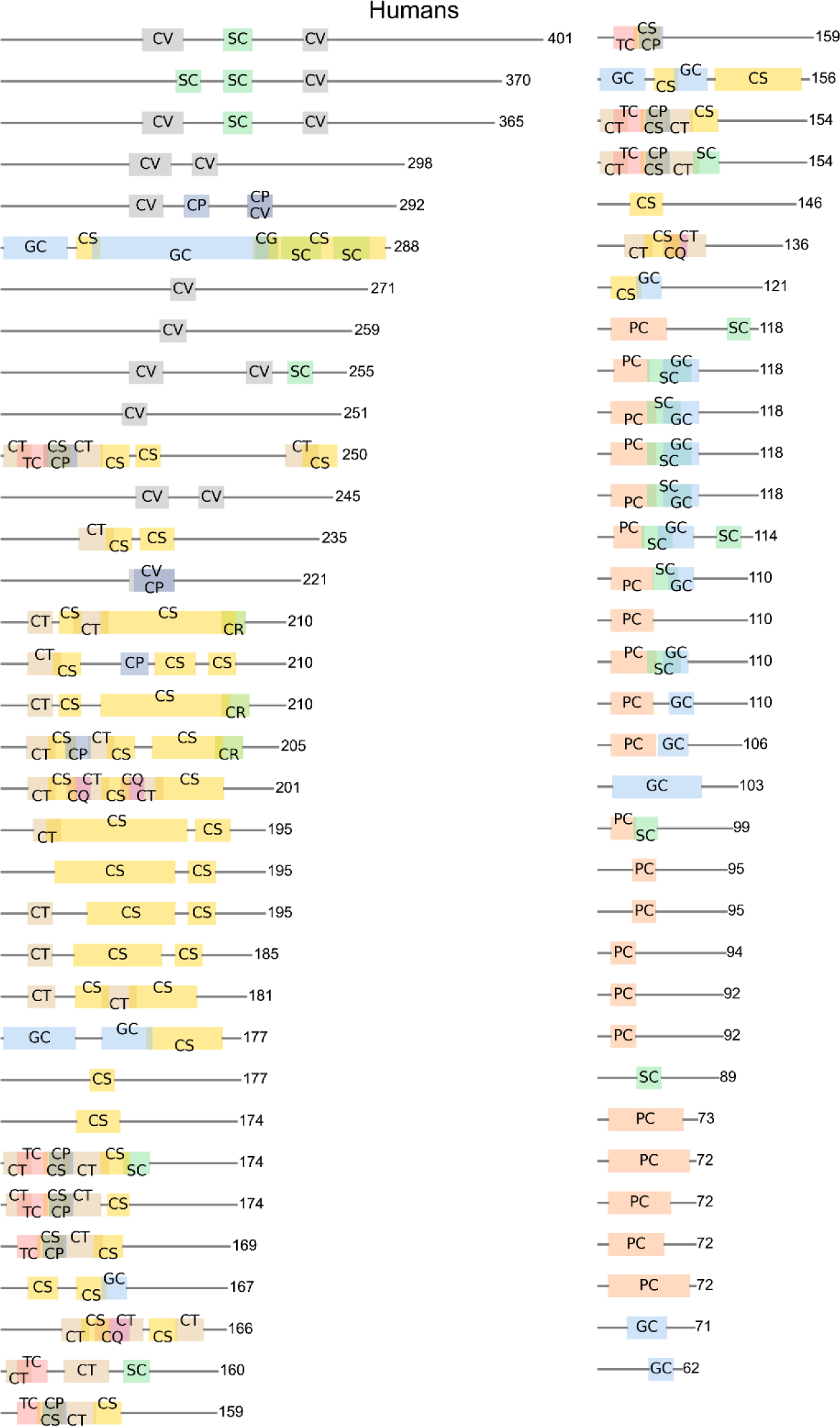
LCD domain layout within human keratin and keratin-associated proteins. Schematic depicting LCD types and their locations within human keratin/keratin-associated proteins. LCD types are indicated by their two-letter abbreviations, and labels are staggered in cases where the LCDs overlap.

**Fig S3.**
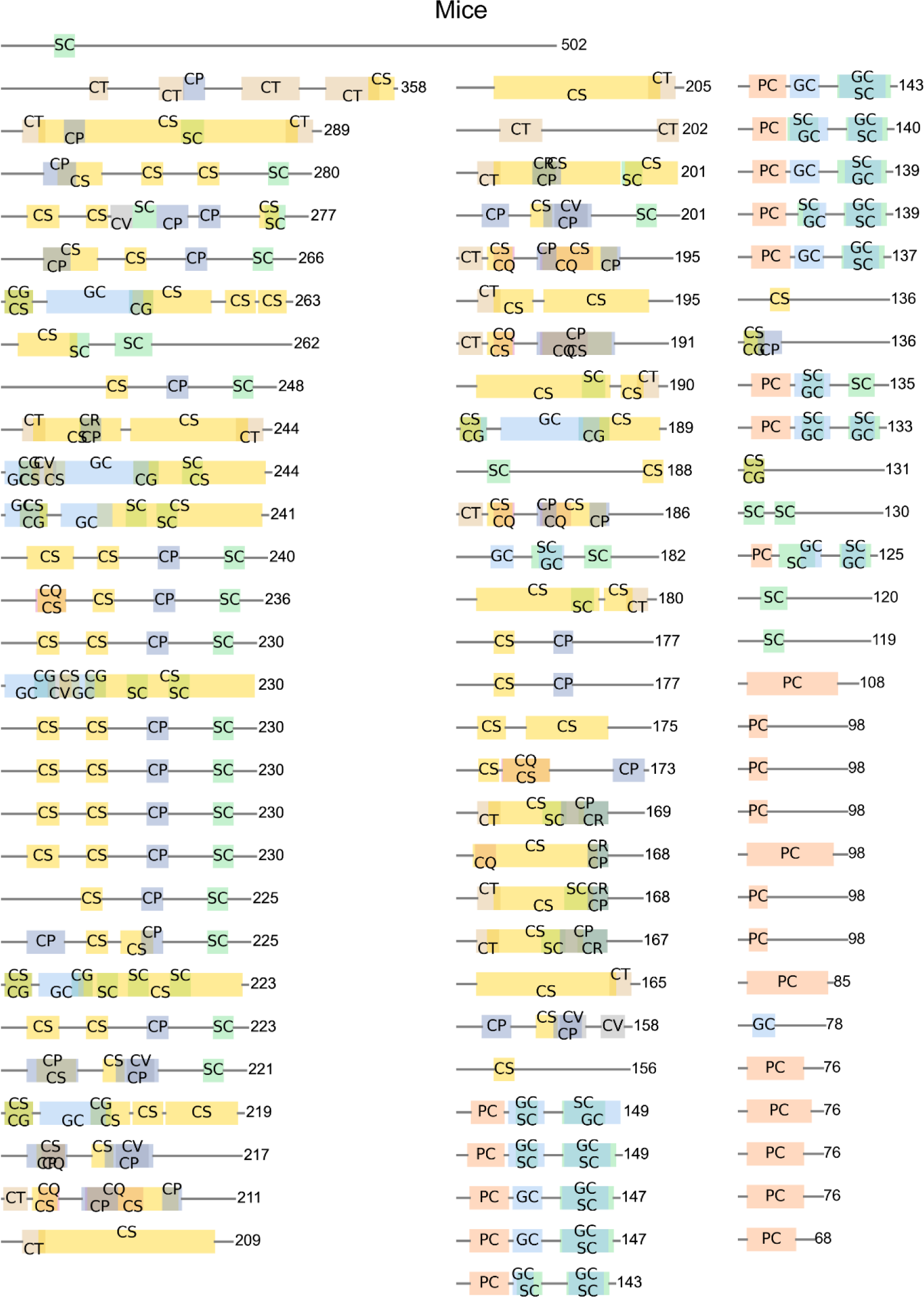
LCD domain layout within mouse keratin and keratin-associated proteins. Same as Fig S2, but for mouse keratin/keratin-associated proteins.

**Fig S4.**
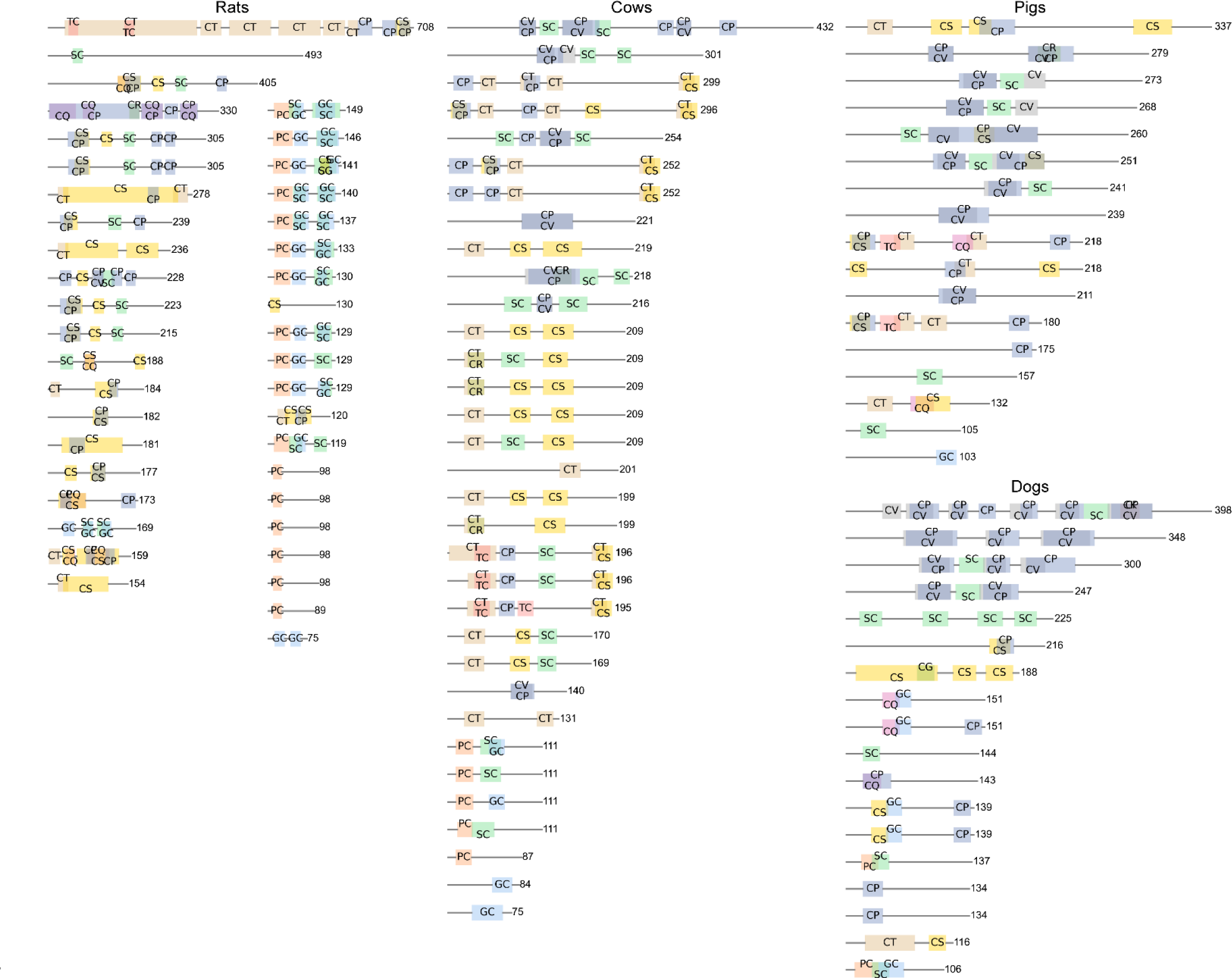
LCD domain layout within rat, cow, pig, and dog keratin and keratin-associated proteins. Same as Fig S2, but for rat, cow, pig, and dog keratin/keratin-associated proteins.

**Fig S5.**
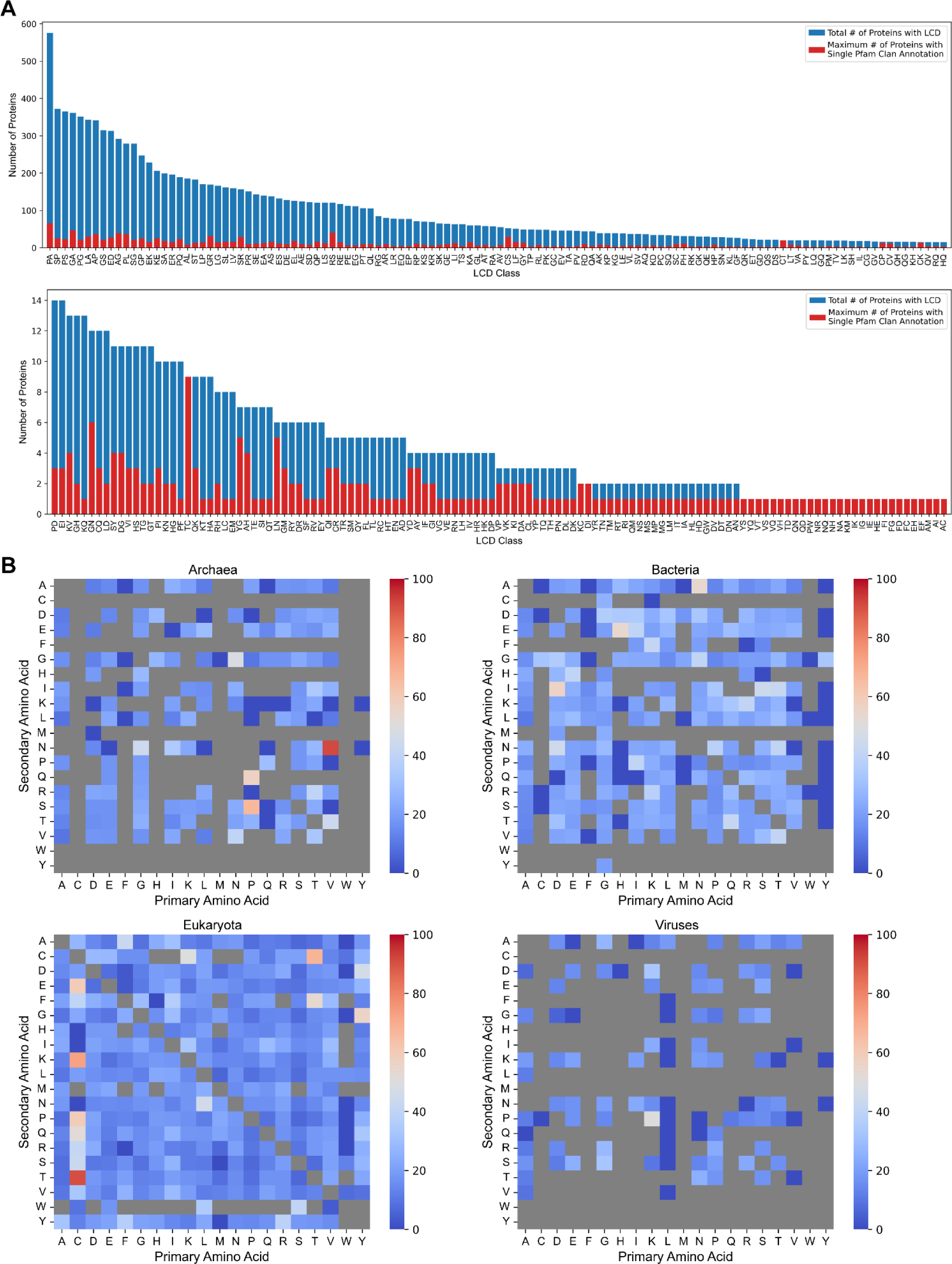
Analysis of Pfam clan annotations associated with each LCD class. (A) Total number of proteins (blue) and the number of proteins associated with a single Pfam clan (red) were calculated for all LCD classes in the human proteome. (B) The average of maximum percentages of proteins associated with each LCD class was calculated across organisms within each domain. A minimum of 5 proteins were required to be included in the calculation to mitigate skewing of percentages by small sample sizes. Grey boxes indicate primary LCD classes and LCD classes for which none of the organisms had ≥5 associated proteins.

**Fig S6.**
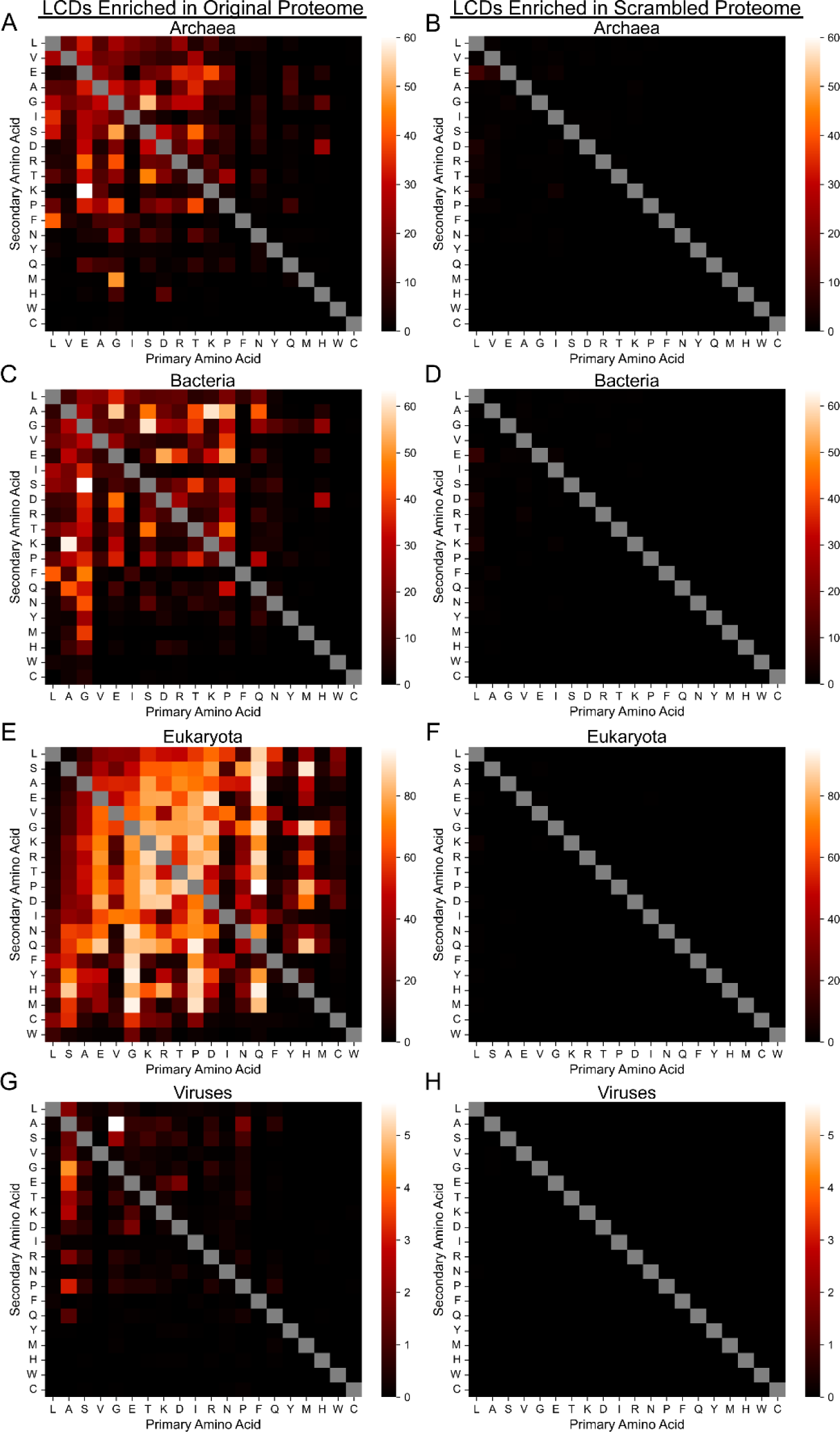
Percentage of organisms with enrichment or depletion of LCDs, regardless of statistical significance. The percentage of organisms with enrichment (positive lnORs) or depletion (negative lnORs) was calculated as described for Fig 4 but without imposing a statistical significance threshold. The panels in the left column (panels A, C, E, and G) indicate the percentage of organisms with LCD enrichment in the original proteome for each LCD class. The panels in the right column (panels B, D, F, and H) indicate the percentage of organisms with LCD depletion in the original proteome for each LCD class. For each pair of heatmaps corresponding to a single domain of life, the heatmap scales are identical to facilitate direct comparison.

**Fig S7.**
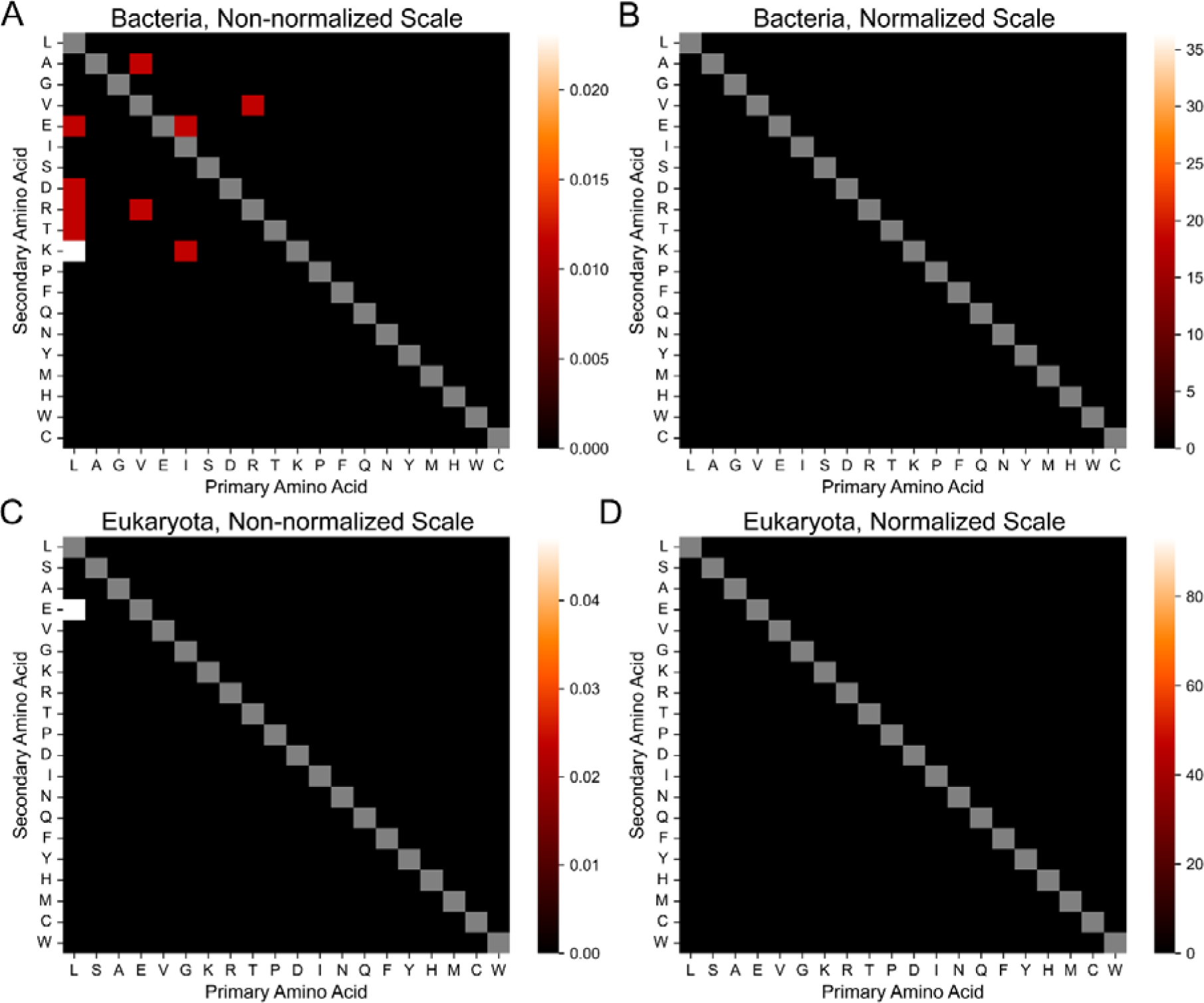
Percentage of organisms with statistically significant depletion of LCDs. The percentage of organisms with statistically significant depletion (negative lnORs) was calculated as described for Fig 4. Heatmaps depict percentages for bacteria (panels A and B) and eukaryotes (panels C and D) only, as all values for archaea and viruses were exactly zero. Panels in the left column depict heatmaps with scales set by the minimum and maximum values within each heatmap. Panels in the right column depict heatmaps with scales identical to those in Fig 4 to facilitate direct comparison.

**Fig S8.**
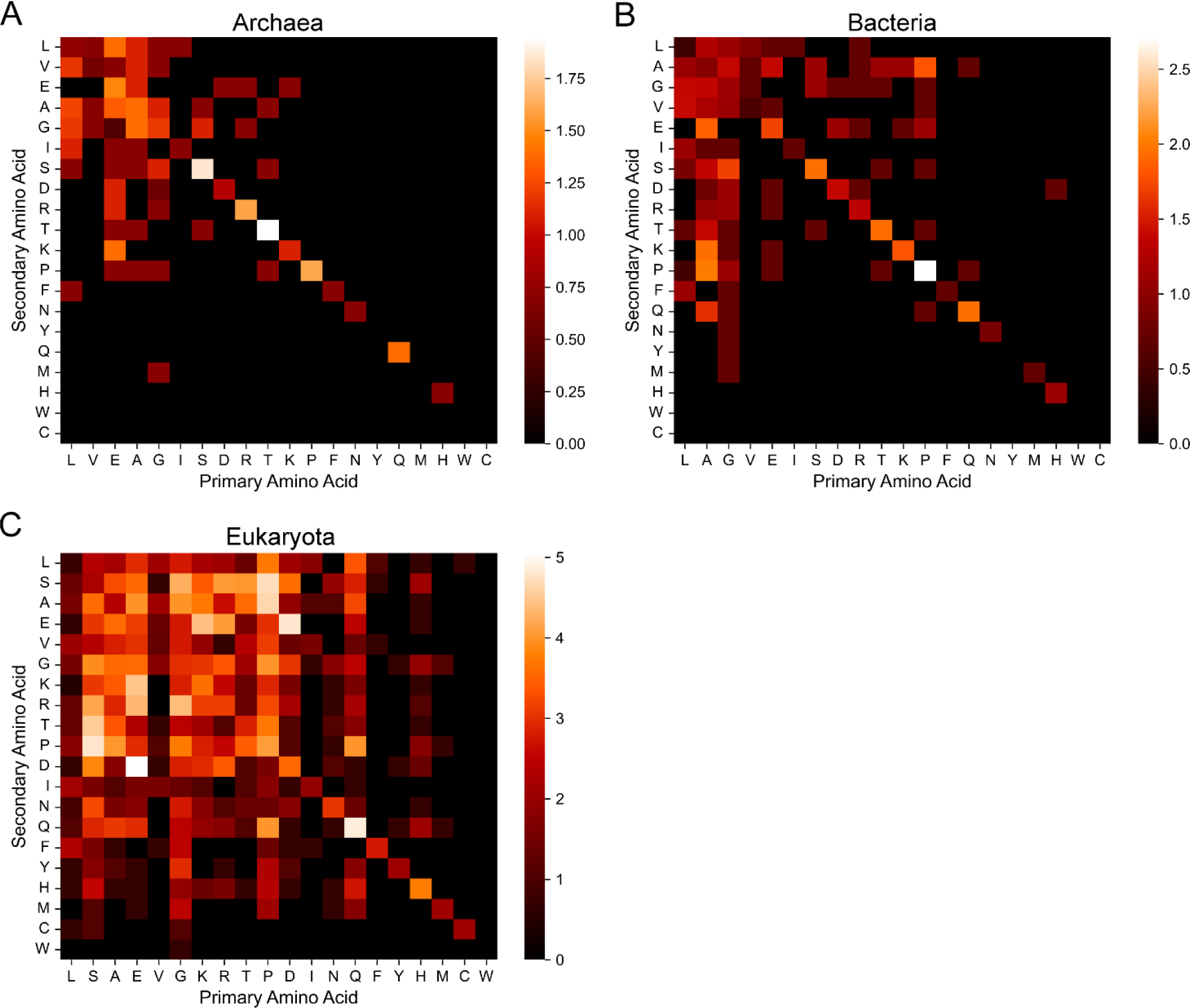
Median degree of LCD enrichment for each LCD class across the domains of life. For each organism, the natural logarithm of the odds ratio (lnOR) for each LCD class was used to quantify the degree of LCD enrichment or depletion in the original proteome relative to a scrambled version of that proteome. Heatmaps depict the median lnOR for each LCD class among archaea (A), bacteria (B), and eukaryotes (C). LCD enrichment data are not shown for viruses since all median lnORs are exactly 0, and no negative median lnOR values were observed for any of the domains of life. The diagonals indicate the median lnOR for each primary LCD class. For LCD classes in which the number of LCDs in either the original or scrambled proteomes were 0, a value of 1 was added to all cells in the contingency table to calculate a biased lnOR (see Methods).

**Fig S9.**
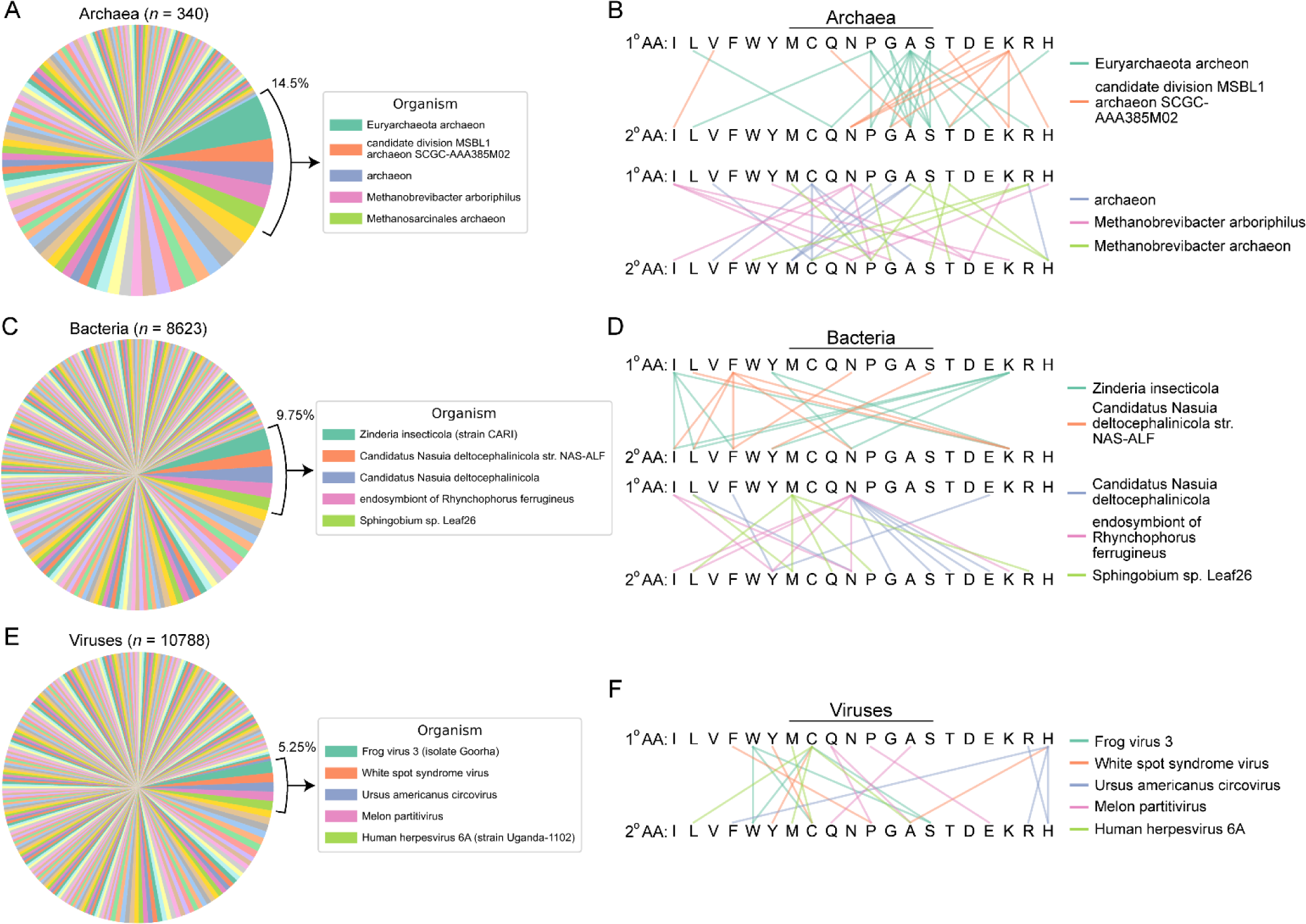
Number of LCD classes assigned to each archaeal, bacterial, or viral organism contributing a maximum per-residue occupancy for at least one LCD class. (A) Pie chart indicating the assignment of LCD classes (400 total) to the archaeal organism achieving the highest per-residue LCD occupancy. Each wedge represents a single organism associated with the overall highest per-residue occupancy observed among bacteria. Wedge size indicates the number of LCD classes for which the single organism corresponding to that wedge achieved the highest per-residue occupancy. (B) Linkage maps indicating the types of LCD classes for which the organism contributed the maximum per-residue occupancy value for eukaryotes. The first row of amino acids in each linkage map indicates the primary amino acid in the LCD class, and lines connected to the second row of amino acids indicate the secondary amino acid in the LCD class. Lines connecting identical amino acids indicate that the organism contributed the maximum per-residue occupancy value for the primary LCD class as a whole (e.g., the W-rich primary LCD class). LCD classes without connecting lines are those for which the organism did not contribute the maximum per-residue occupancy value. Identical analyses were performed for bacteria (C,D) and viruses (E,F). For all pie charts, the top five organisms are indicated in the legend. Out of necessity, the color palette was repeated in each pie chart, though each color cycle represents a different set of organisms.

**Table S1. Number of proteins with LCDs for each LCD class and organism.** Column headers for columns 7-407 indicate the LCD class, with primary LCD classes indicated by a single amino acid and secondary LCD classes indicated by two amino acids. Values indicate the number of proteins in the corresponding organism containing at least one LCD of the indicated LCD class.

**Table S2. Organism-level LCD frequencies among archaea for all LCD classes.** Column headers indicate the primary amino acid in the LCD class, whereas row headers indicate the secondary amino acid in the LCD class. The diagonal (where the primary and secondary amino acid are identical) indicate values for the primary LCD classes. Values indicate the percentage of archaeal organisms containing at least one instance of the indicated LCD class. Columns and rows are sorted by mean amino acid frequency rank across archaeal organisms.

**Table S3. Organism-level LCD frequencies among bacteria for all LCD classes.** Same as Table S2 but with values corresponding to bacterial organisms.

**Table S4. Organism-level LCD frequencies among eukaryotes for all LCD classes.** Same as Table S2 but with values corresponding to eukaryotic organisms.

**Table S5. Organism-level LCD frequencies among viruses for all LCD classes.** Same as Table S2 but with values corresponding to viruses.

**Table S6. Keratin-associated proteins with C-rich LCDs in model organisms.** For each C- rich LCD class containing at least one protein associated with the GO terms “keratin filament” or “keratinization” in the GO term analyses, the UniProt IDs of the corresponding proteins for the indicated LCD class are listed for select model organisms.

**Table S7. GO-term analysis summary for proteins with HQ LCDs in eukaryotes.** For each organism, GO-term analysis was performed on three related sets of proteins: 1) all proteins with an HQ LCD; 2) all proteins with an HQ LCD but excluding those at least 20 residues within a spatially distinct QX, XQ, HX, or XH LCD (where X is any amino acid except H for the QX and XQ classes, and X is any amino acid except Q for the HX and XH classes); and 3) all proteins with an HQ LCD but excluding those with at least 20 residues within a spatially distinct Q-rich primary LCD. None of the proteins had an H-rich primary LCD that was spatially distinct from the HQ domains. To reduce the number of files and file size, only statistically significant GO-terms (Šidák adjusted *p*<0.05) with a minimum depth of 4 in the gene ontology were collected from individual results files.

**Table S8. GO-term analysis summary for proteins with NM, NH, or HL LCDs in eukaryotes.** For each organism with at least 10 proteins containing an NM, NH, or HL LCD (evaluated separately), GO-term analysis was performed on the set of LCD-containing proteins. To reduce the number of files and file size, only statistically significant GO-terms (Šidák adjusted *p*<0.05) were collected from individual results files.

**Table S9. Per-residue-level statistics for the 3 highest scoring organisms across LCD classes for each domain of life.** The first two columns indicate the LCD class and the domain of life, respectively. The last column indicates the average per-residue occupancy for the indicated LCD class within the indicated domain of life. The remaining columns indicate the maximum per-residue occupancy, total number of proteins in the proteome, and total number of amino acids in the proteome for the 3 organisms with the highest per-residue LCD occupancy for the given LCD class and domain of life. These values and the corresponding organism names are ordered from highest-to-lowest per-residue occupancy and are separated by semicolons within each cell.

## References

1. Wootton JC, Federhen S. Statistics of local complexity in amino acid sequences and sequence databases. Comput Chem. 1993;17: 149–163. doi:10.1016/0097-8485(93)85006-X

2. Cascarina SM, Elder MR, Ross ED. Atypical Structural Tendencies Among Low-Complexity Domains in the Protein Data Bank Proteome. PLOS Comput Biol. Public Library of Science; 2020;16. doi:10.1371/journal.pcbi.1007487

3. Kumari B, Kumar R, Kumar M. Low complexity and disordered regions of proteins have different structural and amino acid preferences. Mol Biosyst. 2015;11: 585–594. doi:10.1039/c4mb00425f

4. Cascarina SM, King DC, Osborne Nishimura E, Ross ED. LCD-Composer: an intuitive, composition-centric method enabling the identification and detailed functional mapping of low-complexity domains. NAR Genom Bioinform. Oxford Academic; 2021;3: lqab048. doi:10.1093/nargab/lqab048

5. Harrison PM. Exhaustive assignment of compositional bias reveals universally prevalent biased regions: analysis of functional associations in human and Drosophila. BMC Bioinformatics. 2006;7: 441. doi:10.1186/1471-2105-7-441

6. Yoshizawa T, Nozawa R-S, Jia TZ, Saio T, Mori E. Biological phase separation: cell biology meets biophysics. Biophys Rev. Biophys Rev; 2020;12: 519–539. doi:10.1007/s12551-020-00680-x

7. Dignon GL, Best RB, Mittal J. Biomolecular Phase Separation: From Molecular Driving Forces to Macroscopic Properties. Annu Rev Phys Chem. 2020;71: 53–75. doi:10.1146/annurev-physchem-071819-113553

8. Jalihal AP, Schmidt A, Gao G, Little SR, Pitchiaya S, Walter NG. Hyperosmotic phase separation: Condensates beyond inclusions, granules and organelles. J Biol Chem. Elsevier; 2021;296: 100044. doi:10.1074/jbc.REV120.010899

9. Boeynaems S, Alberti S, Fawzi NL, Mittag T, Polymenidou M, Rousseau F, et al. Protein Phase Separation: A New Phase in Cell Biology. Trends Cell Biol. Elsevier; 2018;28: 420–435. doi:10.1016/J.TCB.2018.02.004

10. Azaldegui CA, Vecchiarelli AG, Biteen JS. The emergence of phase separation as an organizing principle in bacteria. Biophys J. Cell Press; 2021;120: 1123–1138. doi:10.1016/j.bpj.2020.09.023

11. Cinar H, Fetahaj Z, Cinar S, Vernon RM, Chan HS, Winter RHA. Temperature, Hydrostatic Pressure, and Osmolyte Effects on Liquid-Liquid Phase Separation in Protein Condensates: Physical Chemistry and Biological Implications. Chemistry (Easton). Chemistry; 2019;25: 13049–13069. doi:10.1002/chem.201902210

12. Zhang M, Zhu C, Duan Y, Liu T, Liu H, Su C, et al. The intrinsically disordered region from PP2C phosphatases functions as a conserved CO2 sensor. Nat Cell Biol. Nature Publishing Group; 2022;24: 1029–1037. doi:10.1038/s41556-022-00936-6

13. Yang YS, Kato M, Wu X, Litsios A, Sutter BM, Wang Y, et al. Yeast Ataxin-2 Forms an Intracellular Condensate Required for the Inhibition of TORC1 Signaling during Respiratory Growth. Cell. 2019;177: 697–710.e17. doi:10.1016/j.cell.2019.02.043

14. Kato M, Yang YS, Sutter BM, Wang Y, McKnight SL, Tu BP. Redox State Controls Phase Separation of the Yeast Ataxin-2 Protein via Reversible Oxidation of Its Methionine-Rich Low-Complexity Domain. Cell. 2019;177: 711–721.e8. doi:10.1016/j.cell.2019.02.044

15. Oroz J, Félix SS, Cabrita EJ, Laurents D V. Structural transitions in Orb2 prion-like domain relevant for functional aggregation in memory consolidation. J Biol Chem. 2020;295: 18122–18133. doi:10.1074/jbc.RA120.015211

16. Hervas R, Rau MJ, Park Y, Zhang W, Murzin AG, Fitzpatrick JAJ, et al. Cryo-EM structure of a neuronal functional amyloid implicated in memory persistence in Drosophila. Science. American Association for the Advancement of Science; 2020;367: 1230–1234. doi:10.1126/science.aba3526

17. Jung J-H, Barbosa AD, Hutin S, Kumita JR, Gao M, Derwort D, et al. A prion-like domain in ELF3 functions as a thermosensor in Arabidopsis. Nature. Nature Publishing Group; 2020;585: 256–260. doi:10.1038/s41586-020-2644-7

18. Malinovska L, Palm S, Gibson K, Verbavatz J-M, Alberti S. Dictyostelium discoideum has a highly Q/N-rich proteome and shows an unusual resilience to protein aggregation. Proc Natl Acad Sci U S A. 2015;112: E2620–E2629. doi:10.1073/pnas.1504459112

19. Santarriaga S, Petersen A, Ndukwe K, Brandt A, Gerges N, Scaglione JB, et al. The Social Amoeba Dictyostelium discoideum Is Highly Resistant to Polyglutamine Aggregation. J Biol Chem. American Society for Biochemistry and Molecular Biology; 2015;290: 25571–25578. doi:10.1074/jbc.M115.676247

20. Malinovska L, Alberti S. Protein misfolding in Dictyostelium: Using a freak of nature to gain insight into a universal problem. Prion. Taylor & Francis; 2015;9: 339–346. doi:10.1080/19336896.2015.1099799

21. Muralidharan V, Oksman A, Pal P, Lindquist S, Goldberg DE. Plasmodium falciparum heat shock protein 110 stabilizes the asparagine repeat-rich parasite proteome during malarial fevers. Nat Commun. Nature Publishing Group; 2012;3: 1310. doi:10.1038/ncomms2306

22. Wei W, Tan Y, Martinez Rodriguez NR, Yu J, Israelachvili JN, Waite JH. A mussel-derived one component adhesive coacervate. Acta Biomater. Elsevier; 2014;10: 1663– 1670. doi:10.1016/j.actbio.2013.09.007

23. Ahn BK, Das S, Linstadt R, Kaufman Y, Martinez-Rodriguez NR, Mirshafian R, et al. High-performance mussel-inspired adhesives of reduced complexity. Nat Commun. Nature Publishing Group; 2015;6: 8663. doi:10.1038/ncomms9663

24. Valois E, Mirshafian R, Waite JH. Phase-dependent redox insulation in mussel adhesion. Sci Adv. American Association for the Advancement of Science; 2020;6. doi:10.1126/sciadv.aaz6486

25. Cascarina SM, Ross ED. The LCD-Composer Webserver: High-Specificity Identification and Functional Analysis of Low-Complexity Domains in Proteins. Bioinformatics. Bioinformatics; 2022;38: 5446–5448. doi:10.1093/bioinformatics/btac699

26. Cascarina SM, Ross ED. Low-Complexity Domains (LCDs) in UniProt Reference Proteomes. Zenodo. 2023; doi:10.5281/zenodo.8155290

27. Strnad P, Usachov V, Debes C, Gräter F, Parry DAD, Omary MB. Unique amino acid signatures that are evolutionarily conserved distinguish simple-type, epidermal and hair keratins. J Cell Sci. Company of Biologists; 2011;124: 4221–4232. doi:10.1242/jcs.089516

28. Krȩżel A, Maret W. The Bioinorganic Chemistry of Mammalian Metallothioneins. Chem Rev. American Chemical Society; 2021;121: 14594–14648. doi:10.1021/acs.chemrev.1c00371

29. Pande J, Vašák M, Kägi JHR. Interaction of Lysine Residues with the Metal Thiolate Clusters in Metallothionein. Biochemistry. American Chemical Society; 1985;24: 6717–6722. doi:10.1021/bi00344a062

30. Vašák M, McClelland CE, Hill HAO, Kägi JHR. Role of lysine side chains in metallothionein. Experientia. Birkhäuser-Verlag; 1985;41: 30–34. doi:10.1007/BF02005857

31. Jiang L-J, Vašák M, Vallee BL, Maret W. Zinc transfer potentials of the α- and β-clusters of metallothionein are affected by domain interactions in the whole molecule. Proc Natl Acad Sci U S A. The National Academy of Sciences; 2000;97: 2503–2508. doi:10.1073/pnas.97.6.2503

32. Ye B, Maret W, Vallee BL. Zinc metallothionein imported into liver mitochondria modulates respiration. Proc Natl Acad Sci U S A. The National Academy of Sciences; 2001;98: 2317–2322. doi:10.1073/pnas.041619198

33. Cody CW, Huang PC. Metallothionein Detoxification Function Is Impaired by Replacement of Both Conserved Lysines with Glutamines in the Hinge between the Two Domains. Biochemistry. American Chemical Society; 1993;32: 5127–5131. doi:10.1021/bi00070a022

34. Urano F, Calfon M, Yoneda T, Yun C, Kiraly M, Clark SG, et al. A survival pathway for Caenorhabditis elegans with a blocked unfolded protein response. J Cell Biol. J Cell Biol; 2002;158: 639–646. doi:10.1083/jcb.200203086

35. Kamal M, Tokmakjian L, Knox J, Mastrangelo P, Ji J, Cai H, et al. A spatiotemporal reconstruction of the C. elegans pharyngeal cuticle reveals a structure rich in phase-separating proteins. eLife. eLife Sciences Publications Ltd; 2022;11: e79396. doi:10.7554/elife.79396

36. Eckhart L, Ehrlich F. Evolution of trichocyte keratins. Adv Exp Med Biol. Springer New York LLC; 2018;1054: 33–45. doi:10.1007/978-981-10-8195-8_4

37. Pizzi E, Frontali C. Low-Complexity Regions in Plasmodium falciparum Proteins. Genome Res. Cold Spring Harbor Laboratory Press; 2001;11: 218–229. doi:10.1101/gr.152201

38. Cascarina SM, Ross ED. Expansion and functional analysis of the SR-related protein family across the domains of life. RNA. Cold Spring Harbor Laboratory Press; 2022;28: 1298–1314. doi:10.1261/rna.079170.122

39. Thandapani P, O’Connor TR, Bailey TL, Richard S. Defining the RGG/RG Motif. Molecular Cell. 2013. pp. 613–623. doi:10.1016/j.molcel.2013.05.021

40. Süveges D, Gáspári Z, Tóth G, Nyitray L. Charged single α-helix: A versatile protein structural motif. Proteins Struct Funct Bioinform. 2009;74: 905–916. doi:10.1002/prot.22183

41. Sivaramakrishnan S, Spink BJ, Sim AYL, Doniach S, Spudich JA. Dynamic charge interactions create surprising rigidity in the ER/K α-helical protein motif. Proc Natl Acad Sci U S A. National Academy of Sciences; 2008;105: 13356–13361. doi:10.1073/pnas.0806256105

42. Chou C-C, Wang AHJ. Structural D/E-rich repeats play multiple roles especially in gene regulation through DNA/RNA mimicry. Mol Biosyst. 2015;11: 2144–2151. doi:10.1039/c5mb00206k

43. Lee C-H, Shih Y-P, Ho M-R, Wang AHJ. The C-terminal D/E-rich domain of MBD3 is a putative Z-DNA mimic that competes for Zα DNA-binding activity. Nucleic Acids Res. Oxford Academic; 2018;46: 11806–11821. doi:10.1093/nar/gky933

44. Voolstra CR, Aranda M, Zhan Y, Dekker J. Symbiodinium microadriaticum (coral microalgal endosymbiont). Trends Genet. Elsevier Ltd; 2021;37: 1044–1045. doi:10.1016/j.tig.2021.08.008

45. Nand A, Zhan Y, Salazar OR, Aranda M, Voolstra CR, Dekker J. Genetic and spatial organization of the unusual chromosomes of the dinoflagellate Symbiodinium microadriaticum. Nat Genet. Nature Publishing Group; 2021;53: 618–629. doi:10.1038/s41588-021-00841-y

46. Cascarina SM, Ross ED. Proteome-scale relationships between local amino acid composition and protein fates and functions. PLOS Comput Biol. Public Library of Science; 2018;14: e1006256. doi:10.1371/journal.pcbi.1006256

47. Radó-Trilla N, Albà M. Dissecting the role of low-complexity regions in the evolution of vertebrate proteins. BMC Evol Biol. 2012;12: 155. doi:10.1186/1471-2148-12-155

48. Sim KL, Creamer TP. Abundance and Distributions of Eukaryote Protein Simple Sequences. Mol Cell Proteomics. 2002;1: 983–995. doi:10.1074/mcp.M200032-MCP200

49. Albà MM, Guigó R. Comparative analysis of amino acid repeats in rodents and humans. Genome Res. 2004;14: 549–554. doi:10.1101/gr.1925704

50. Faux NG, Bottomley SP, Lesk AM, Irving JA, Morrison JR, De La Banda MG, et al. Functional insights from the distribution and role of homopeptide repeat-containing proteins. Genome Res. 2005;15: 537–551. doi:10.1101/gr.3096505

51. Gutierrez JI, Brittingham GP, Karadeniz Y, Tran KD, Dutta A, Holehouse AS, et al. SWI/SNF senses carbon starvation with a pH-sensitive low-complexity sequence. eLife. eLife Sciences Publications Ltd; 2022;11: e70344. doi:10.7554/eLife.70344

52. Corbet GA, Wheeler JR, Parker R, Weskamp K. TDP43 ribonucleoprotein granules: physiologic function to pathologic aggregates. RNA Biol. Taylor & Francis; 2021;18: 128–138. doi:10.1080/15476286.2021.1963099

53. Cui M, Wang X, An B, Zhang C, Gui X, Li K, et al. Exploiting mammalian low-complexity domains for liquid-liquid phase separation–driven underwater adhesive coatings. Sci Adv. American Association for the Advancement of Science; 2019;5: eaax3155. doi:10.1126/sciadv.aax3155

54. Klopfenstein D V., Zhang L, Pedersen BS, Ramírez F, Vesztrocy AW, Naldi A, et al. GOATOOLS: A Python library for Gene Ontology analyses. Sci Rep. 2018;8. doi:10.1038/s41598-018-28948-z

